# Blood-brain-barrier permeable fluorescent astrocyte probes

**DOI:** 10.1101/2025.07.10.664185

**Authors:** Nipuni Gunawardhana, Danielle A. Cervasio, Shangrila Singh, Oshini Haputhanthrige-Don, Scott T. Laughlin

## Abstract

The brain’s astrocytes play pivotal roles in brain homeostasis, but their contributions to neural circuit function are only beginning to be explored. In order to study astrocytes in their native environment, the field needs imaging probes that are cell-specific, non-toxic, and complementary to existing genetic techniques. We recently described a panel of cationic fluorophores that selectively label rodent and zebrafish astrocytes at low concentrations and short incubation times. Here, we expand on those first-generation astrocyte probes with a new class of brighter and more photostable dyes that allow for imaging across various wavelengths. We demonstrate these probes’ subcellular localization to the mitochondria and ability to cross the blood-brain barrier in zebrafish. This new class of astrocyte markers will aid researchers in visualizing and identifying subpopulations of astrocytes in both rodent and zebrafish model systems.

---

Astrocytes are the most abundant glial cell-type in the brain and have critical functions in every layer from the olfactory bulb to the cortices and midbrain.^1^ They are active players in diverse brain processes like hormone metabolism, ion buffering, and neuronal signaling.^2,3^ Given their important roles in brain function, it is not surprising that astrocytes are implicated in many diseases, including Alzheimer’s, Parkinson’s, glioblastoma, stroke, seizure, and depression.^4^ Accordingly, it is essential to be able to visualize and study these cells in their native environments so we can both better understand their biology and develop therapeutic interventions for astrocyte-associated disease.

While there exist a handful of astrocyte-specific genes whose promoters are utilized for genetic targeting and visualization (*e*.*g*., GFAP^5^, AldH1L1^6^, S100β^7,8^), the process of creating transgenic organisms is time consuming and costly, and promoter-driven fluorescent reporters often fail to act as pan-astrocyte markers^6^ due to the heterogeneity of the astrocyte cell-type. Indeed, recent studies are beginning to uncover that astrocytes vary greatly between and within brain regions, expressing different genetic markers and performing unique functions.^9,10^ In addition to the genetic markers used for studying astrocytes, researchers have recently explored the use of small molecule probes for astrocytes. By utilizing small molecule astrocyte-specific fluorophores like Sulforhodamine 101^11,12^, Ala-Lys-Coumarin^13^, Cy-BASHY^14^, Rhodamine Q-MP^15^, and Rhodamine B-MP^16,17^ researchers can complement the use of transgenic animals, image across a spectrum of visible and infrared wavelengths, and probe astrocytes *in vivo*.

We recently described a library of cationic dyes that leverage a pyridinium moiety to achieve astrocyte-specific labeling via astrocyte-resident organic cation transporters.^16,17^ Diverse fluorophores including rhodamines and cyanines tagged with the pyridinium group produced strong labeling of mouse, rat, and zebrafish astrocytes, but were excluded from other glia and neurons.^15,16^ Herein, we report next generation astrocyte probes based on the brighter, more photostable Janelia Fluor® fluorophores^18–20^ combined with an added pyridinium to drive astrocyte selectivity. We demonstrate the astrocyte-specificity of these fluorophores in cell culture and *in vivo*. We report labeling in cell culture and mitochondrial colocalization after only 20 minutes of incubation at low micromolar concentrations. Additionally, we demonstrate these probes’ ability to cross the blood brain barrier (BBB) in zebrafish in order to label their astrocyte targets after intravenous injection. The versatility in wavelength and structural modularity of this approach has the potential to uncover information regarding the heterogeneity of cells classified as astrocytes and provide a toolkit for efficient and easy live imaging without the need for specialized microscopy or invasive delivery methods.

We chose to modify three Janelia Fluor® probes with excitation wavelengths ranging from 503 to 585 nm to demonstrate targeted labeling of astrocytes using a subset of these next generation fluorophores. Following previously reported synthetic methods, we began with commercially available fluorescein for Janelia Fluor® 549 and 503.^18,21^ We then appended the methylpyridinium moiety that is responsible for astrocyte targeting via HATU amide coupling in DMF to obtain 549MP, ***4a***, and 503MP, **5*a*** in 29% and 19% yield, respectively (**Figure 1A**). For the synthesis of 585MP, ***6a***, we followed previously reported synthetic methods to obtain the triflated carbofluorescein backbone over 8 steps.^18^ Our attempts at HATU coupling with the latter probe proved unsuccessful, probably due to its near-zero L-Z equilibrium constant, favoring the lactone-form and therefore limiting access to the carboxylic acid as a coupling handle.^19^ Structurally, these fluorescent probes differ either in their xanthene substituent or in their appended nitrogen-containing groups on the xanthene ring from our previously reported probe, Rhodamine B methylpyridinium (Rh_B_MP)^16^. The reported photophysical properties of the second-generation parent fluorophores, JF503, 549, and 585 are superior in comparison to the parent fluorophore of our first probe, Rhodamine B, in their quantum yield and extinction coefficients^19^, which is what inspired us to make these fluorophores amenable to live-imaging in astrocytes.

**Figure 1.**
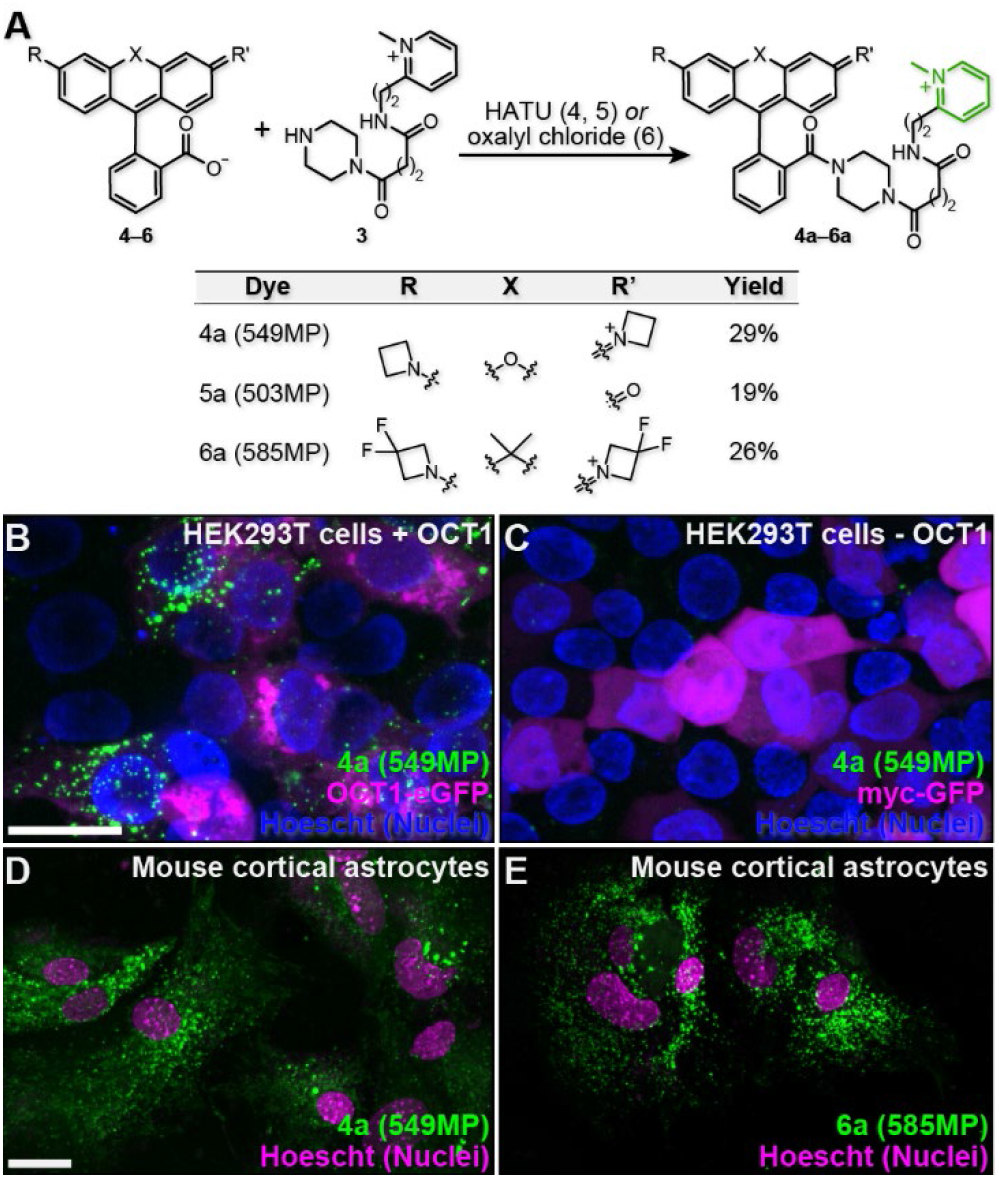
Synthesis and cellular-specific uptake of methyl-pyridinium-modified fluorophores. (**A**) General synthetic scheme for compounds **4a** (549MP), **5a** (503MP), and **6a** (585MP) using HATU coupling or oxalyl chloride, respectively. (**B**–**C**) Compound **4a** labels HEK293T cells expressing OCT1 (***B***), but not cells expressing control plasmid myc-eGFP (***C***) after 20 minutes of bathing at 1 μM. (**D**–**E**) Compounds **4a** (***D***) and **6a** (***E***) label primary mouse cortical astrocytes after 20 mins of bathing with 1 μM. Scale = 50 μm (B), 25 μm (D).

We sought to analyze the probes’ performance in cell culture considering their similar structure, yet superior quantum yield, extinction coefficient, and varying excitation/emission spectra which would make for a diverse tool-kit of brighter, more photostable^19,20^ and spectrally-diverse astrocyte-targeted probes. We first evaluated **4a** (549MP) in HEK293T cells with or without the overexpression of Organic Cation Transporter 1 (OCT1), which our previous studies have shown is sufficient for cellular uptake of the molecules^16,17^. HEK293T cells transfected with OCT1 were successfully labeled with ***4a*** (**Figure 1B** and **S1**), whereas HEK293T cells transfected with a GFP control did not take up the compound (**Figure 1C** and **S1**), suggesting that it is not cell permeable and is able to transit the OCT1 like its predecessors. We next evaluated ***4a****–****6a*** for their ability to label primary astrocytes in culture. In these experiments, ***4a*** and ***6a*** successfully labeled astrocytes in primary cultures derived from mouse cortex at a working concentration of 1 µM (**Figure 1D, E** and **S2**). Despite its structural similarities, ***5a*** did not show either specific or diffuse labeling in astrocytes, HEK293T cells, or primary cortical culture suggesting that it is neither astrocyte-targeted nor cell permeable (**Figure S2 I**–**J**). We have demonstrated previously that the presence of a +2 charge on the cargo molecule results in more robust targeting of astrocytes^16,17^, which might explain the inability of the rhodol, ***5a***, to label these cells due to its overall +1 charge. On the other hand, ***6a*** shows labeling of astrocytes while bearing a +2 charge and two 3,3-difluoroazetidine moieties at the xanthene ring, albeit less robust than the flagship ***4a*** (**Figure 1E** and **S2 E**–**F**). The electronegative fluorine atoms may change the distribution of electron density, which may decrease uptake by the organic cation transporter responsible for astrocyte labeling.

Next, we evaluated the cytotoxicity of compounds ***4a, 5a***, and ***6a*** using the colorimetric MTT cell viability assay. At a concentration of 1 μM and exposure time of 20 min, the conditions we employ for cultured astrocyte labeling, we observed 90–100% cell viability for all fluorophores in HEK293T cells expressing OCT1 (**Figure S3**).^16^ In the most extreme case, viability in the MTT assay drops to approximately 85% for ***6a***-treated cells when we increase the concentration to 10 μM, 10-fold above the concentrations used for cell labeling, suggesting the probes have minimal toxic effects in mammalian cells (**Figure S3 A**).

We next sought to evaluate the subcellular distribution of compounds ***4a*** and ***6a*** in labeled cells. In previous reports with Rh_B_MP, and here with compounds ***4a*** and ***6a*** (**Figure 1D, E**, and **S2**), we observed a punctate labeling pattern in primary mouse and zebrafish astrocytes^16^. We also observed a similar subcellular distribution in HEK293T and HeLa cells transfected with OCT1 (**Figure 1B, C**, and **S1**). This punctate labeling suggested organelle-specific rather than non-specific or cytosolic subcellular localization. We hypothesized that punctate labeling was mitochondrial, since the pattern is consistent with mitochondrial labeling in cultured cells and there are several small molecule probes for mitochondria that possess a positive charge (Rhodamine 123 and.tetramethylrosamine^22^). To evaluate mitochondrial labeling, we performed a colocalization experiment with MitoTracker® and compounds ***4a*** and ***6a***. After labeling mouse primary cortical astrocytes with 1 µM of the compounds for 20 min, we observed a clear colocalization of ***4a*** and our first-generation probe, Rh_B_MP with the mitochondrial dye, MitoTracker® via confocal microscopy (**Figure 2** and **S4**). This result suggests that fluorophores of varying excitation/emission patterns can be modified in order to target the mitochondria of astrocytes. Both ***4a*** and our first-generation probe, Rh_B_MP strongly label the mitochondria of mouse cortical astrocytes, while ***6a*** does not appear as robust (**Figure 2** and **S4**). The percentage of pixels of ***4a, 6a***, and Rh_B_MP that colocalize with MitoTracker® are 90%, 95%, and 93%, respectively, as calculated by the Mander’s colocalization coefficient (MCC) (**Figure S4**)^23,24^. Moving forward, we decided to focus our efforts on the flagship molecule ***4a*** since it produced the most robust labeling of astrocytes and their mitochondria. Taken together, these probes can be utilized for imaging studies where transgenic organisms may not be available or where visualization of both astrocytes and their mitochondria is desired. These results also give insight into the types of molecules that mitochondria are likely to take up, and imply similar modifications researchers might make in order to target other small molecule cargo to astrocytes.

**Figure 2.**
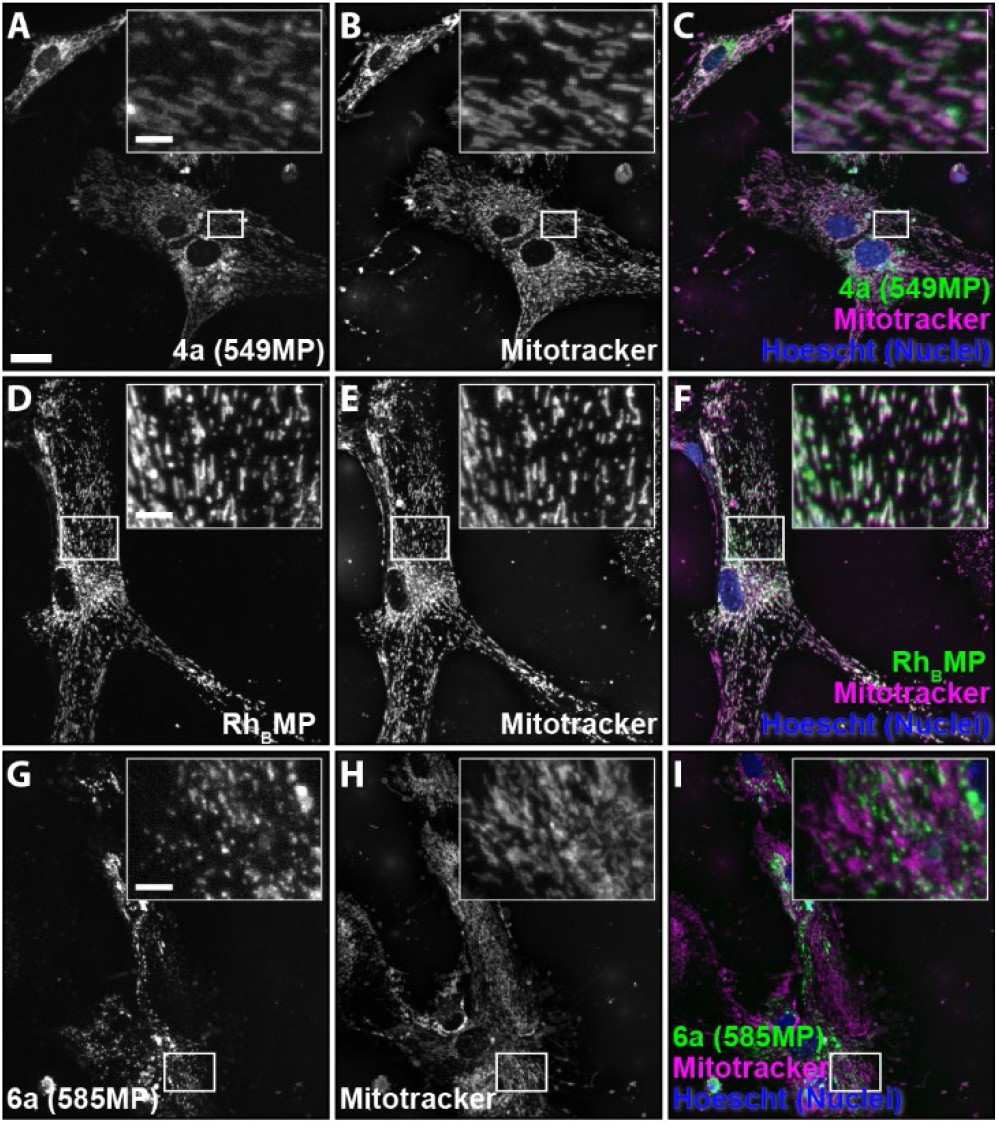
Fluorescent probes colocalize to mitochondria in mouse primary cortical astrocytes. After 20 mins incubation at 1 μM, **4a** (**A**–**C**), Rh_B_MP (**D**–**F**), and **6a** (**G**–**I**) colocalize with MitoTracker in mouse primary cortical astrocytes. Scale = 25 μm.

After confirming that the dye was astrocyte-specific and observing mitochondrial-targeted labeling, we sought to move *in vivo* to assay its performance in a whole organism model system. Zebrafish are frequently used in brainimaging studies due to their ability to produce a large number of embryos quickly and their transparency in larval stages after exposure to the drug N-Phenyl-2-thiourea (PTU). It is well-known that zebrafish contain cells homologous to mammalian astrocytes that express both GFAP and regulate neuronal signaling.^25–27^ Upon direct intracerebroventricular injections of our flagship molecule, ***4a***, we see clear colocalization with the GFAP+ astrocytes of 3–5 days post fertilization (dpf) larval zebrafish, corroborating our cell culture results (**Figure 3** and **S5**). Importantly, no overlap is seen between the probe and neurons, oligodendrocytes, or microglia in this whole-organism system (**Figure 3H–J** and **S5**), suggesting that ***4a*** is indeed astrocyte specific in both mammalian cell culture and in the larval zebrafish brain.

**Figure 3.**
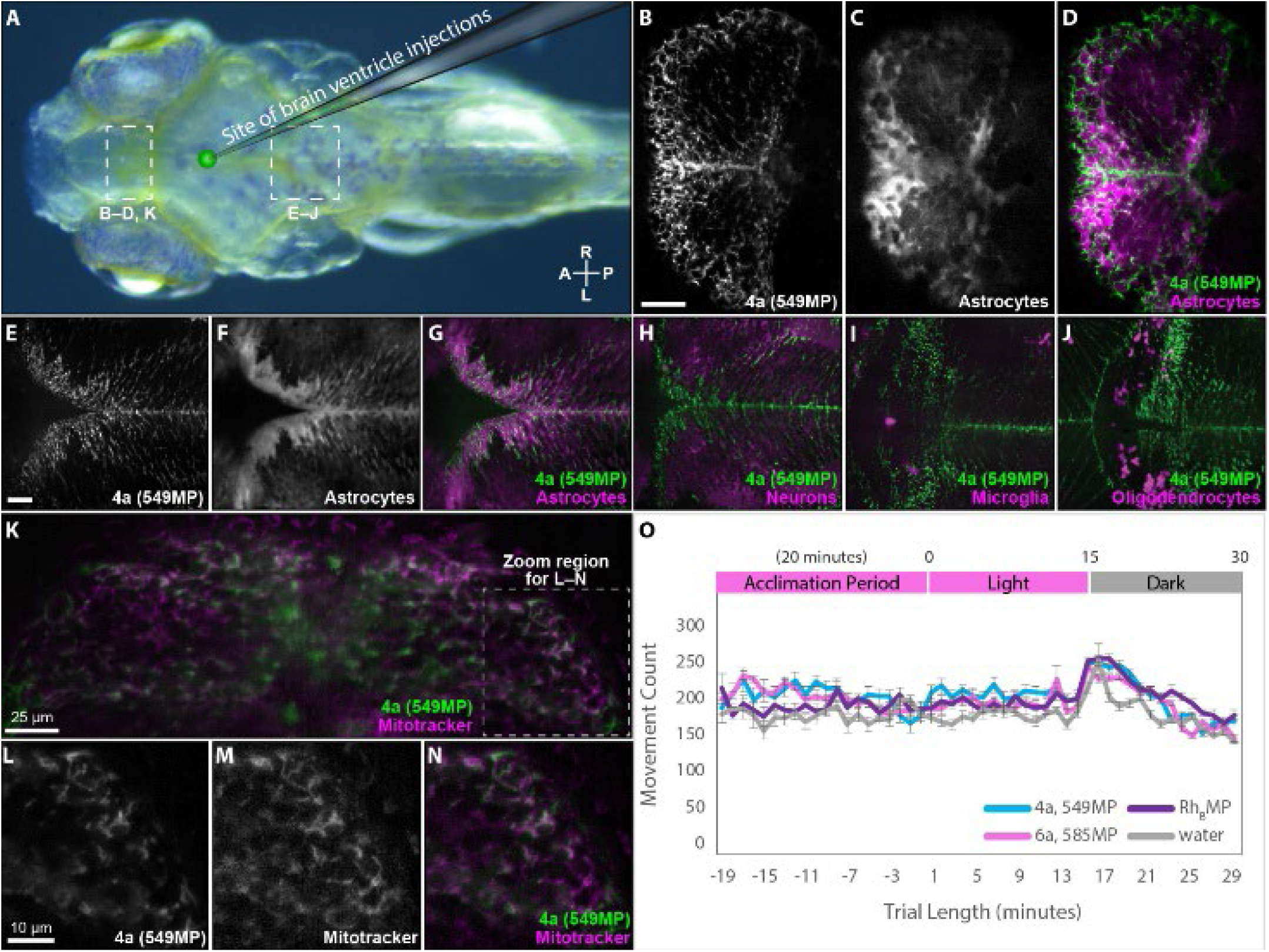
Injection of flagship molecule 4a (549MP) directly into the brain ventricle of 3-5 dpf larval zebrafish results in astrocyte uptake and mitochondrial colocalization. (A) Schematic of a left-facing zebrafish indicating site of ventricle injection (2.3 nL of 100 μM solution). (B–D) Grayscale image of 4a distribution (B), GFAP+ astrocytes (C), and pseudo-colored overlay of both channels (D) showing colocalization between 4a and astrocytes in the telencephalon. (E–G) Grayscale image of 4a labeling (E), GFAP+ astrocytes (F), and pseudo-colored overlay (G) of both channels showing overlap in the hindbrain region. Colocalization is not seen between neurons (H), microglia (I), or oligodendrocytes (J) in the hindbrain. (K) After co-injection of both 4a (600μM) and MitoTracker (400μM), colocalization is seen in the telencephalon. (L–N) Inset from (K) showing 4a labeling (L), MitoTracker labeling (M), and a pseudo-colored overlay (N), indicating overlap of the two fluorophores. (O) Spontaneous and photic-evoked larval swim behavior assay. Line graph showing average movement count of 6 dpf larval zebrafish at 18 hpi of 4a (n = 10), 6a (n = 10), Rh_B_MP (n = 10), and a water control (n = 10). Scale = 25 μm (B, E).

To assay for high level behavior effects of these molecules, we used a spontaneous and light-evoked locomotion paradigm where spontaneous movement is monitored in zebrafish larvae at 6 dpf. Larvae acclimate to a behavior chamber for 20 mins in the light and are then recorded for 15 mins of spontaneous movement followed by removal of illumination, which elicits a visual motor response, a stereotypical zebrafish behavior.^28,29^ This allows for the quantitative assessment of the molecules’ effects on normal zebrafish behavior. All larvae exhibit the typical visual motor response, demonstrated by a temporary increase in movement at 15 mins when light switches to dark (**Figure 3O**). We see no significant difference in movement count, duration, and distance between larvae injected with our probes versus a water control at 18 hours post-injection (hpi) in the acclimation and dark periods (**Figure S5 Q, S**, and **S6 B, D, F, H**). We do see a small increase in movement count during the light-period, between ***4a*** and control (**Figure S5 R**) in addition to increases in the duration of movements during the light period between all probes and the control (**Figure S6 C**). Taken together, fish injected with the fluorescent probes do not show deficits in their locomotor behavior. Additionally, we see no significant difference in additional parameters including distance traveled (**Figure S6 E**–**H**), swim length, and swim speed (**Figure S7**), suggesting that fish injected with our probes do not show behavioral deficits in their overall movement patterns.

Since we had seen mitochondrial colocalization (**Figure 2**) in primary cortical astrocytes, we sought to assay the probes cellular compartmentalization *in vivo*, as well. Accordingly, when both MitoTracker and ***4a*** are co-injected into wild-type (WT-AB) fish, strong colocalization is seen in the telencephalon, optic tectum, and hindbrain regions (**Figure 3K** and **S9 A**–**F**). Furthermore, when MitoTracker, alone, is injected into GFAP-GFP+ fish, we see that it does indeed label GFAP+ cells, of course non-exclusively, since it is meant to label all cells’ mitochondria (**Figure S9 S**–**X**). Undeterred by our unexpected *in vitro* results with ***5a*** and ***6a***, we decided to assess their performance *in vivo*. The rhodol ***5a*** produces labeling (**Figure S8 B**–**C**) and mitochondrial colocalization (**Figure S9 M**–**O**), especially in the telencephalon and hindbrain, albeit less robust and at a higher injection concentration of 1 mM. Moreover, ***6a*** labels astrocytes (**Figure S8 D**–**I**) over neurons and other glial cells (**Figure S8 J**–**R**) after a 1 mM injection, and is also phagocytosed by microglia as demonstrated by its engulfment by cells marked by GFP under control of the mpeg1 microglial-specific-promoter (**Figure S8 M**–**O**). Moreover, we show that ***6a*** colocalizes with mitochondria in hindbrain (**Figure S9 P**–**R**) and our first-generation probe, Rh_B_MP, exhibits strong mitochondrial colocalization in both telencephalon (**Figure S9 G**–**I**) and optic tectum (**Figure S9 J**–**L**). Taken together, these results suggest that our molecules specifically label the mitochondria of astrocytes in zebrafish in addition to mammalian cells in culture.

We next sought to evaluate these astrocyte-targeted fluorophores’ blood-brain barrier (BBB) permeability by injecting them systemically through the pericardial vein and assessing labeling in the CNS. Zebrafish are the smallest vertebrate model with a functional BBB and studies suggest histological and ultrastructural similarities between rodent and zebrafish BBB, making zebrafish a good model for predicting small molecule BBB permeability.^30,31^ The zebrafish BBB begins functioning as early as 2.5 dpf, contains mature endothelial cells, pericytes, and contacting glia by 5 dpf, and a more complex double-layer vessel structure by 10 dpf.^30,32,33^ We injected 2.3 nL of a 1 mM solution of ***4a****–****6a*** and Rh_B_MP, which resulted in rapid distribution of the compounds throughout the vasculature of 7 dpf larval zebrafish (**Supplemental Movie 1**). After allowing the fish to recover for 3 hours we observed robust labeling of ***4a*** in the brain, indicating the molecules’ ability to cross the BBB in this whole organism system, despite its size and charge (**Figure 4B–D** and **S10 A**–**F**). Additionally, Rh_B_MP, ***5a***, and ***6a*** are able to cross the BBB to label their astrocyte counterparts (**Figure 4E–H** and **S10 G**–**S**). Conversely, the unconjugated JF549^®^ and structurally similar MitoTracker^®^ Red FM are both unable to label the brain at the same concentration of 1 mM, showing the BBB is indeed intact, for similarly sized molecules (**Figure 4I–J** and **S10 T**–**X**). This suggests that systemic delivery of these fluorophores might be a reasonable, minimally-invasive route of administration when moving forward into mammals like rodents in order to visualize astrocytes for further live studies since they are able to cross the BBB. This also raises the question of if chemical modification with the methyl-pyridinium moiety is sufficient to permit uptake of other small molecule cargo across the BBB which could be useful for targeted delivery of drugs, for example, to the brain.

**Figure 4.**
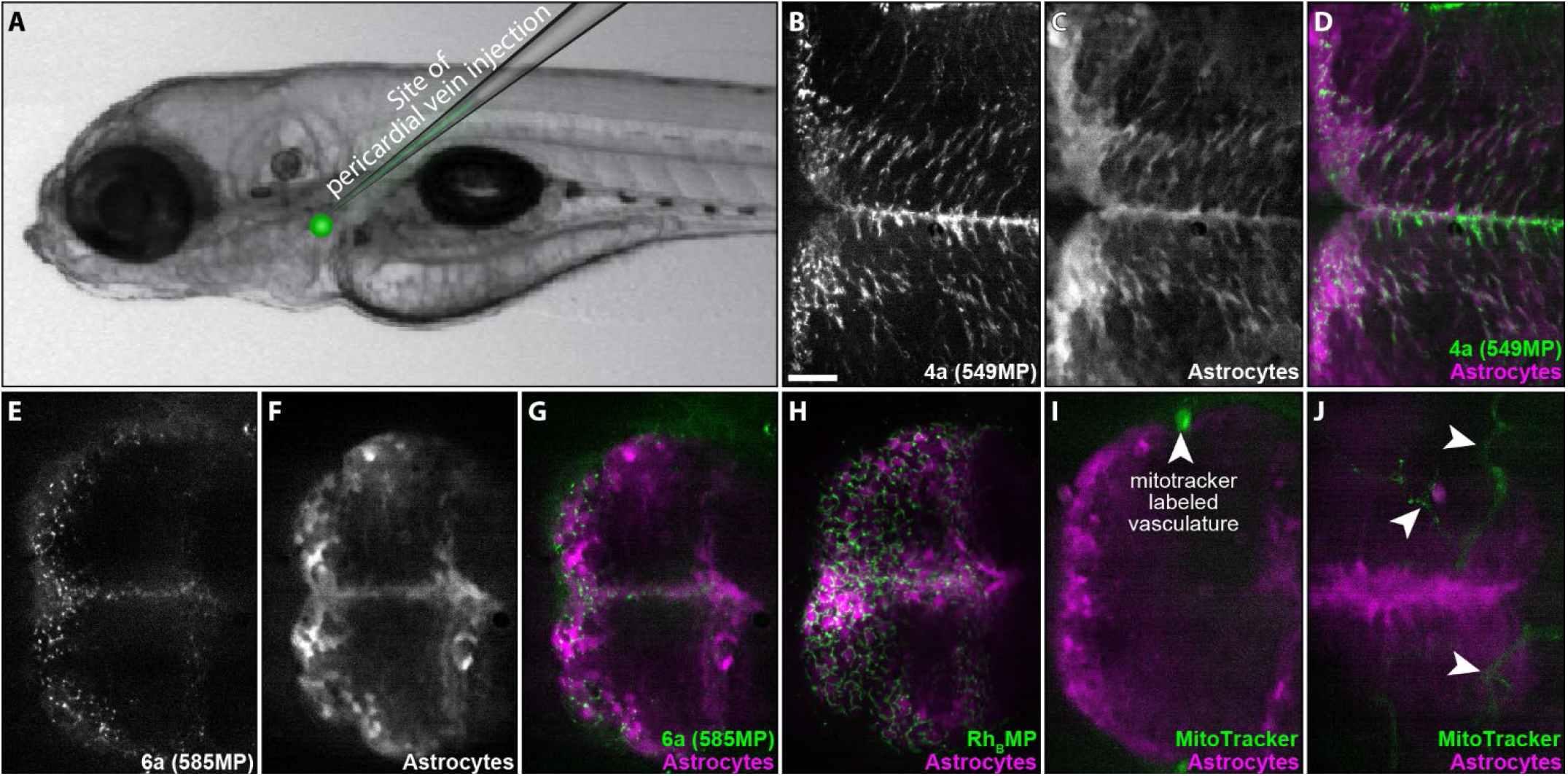
Fluorescent probes cross the BBB and label astrocytes after systemic introduction into 7 dpf GFAP-GFP zebrafish. (**A**) Schematic showing injection location of the fluorophores at 2.3 nL of a 1 mM solution. (**B**–**D**) Crossing of the BBB and colocalization of **4a** (**B**) with astrocytes (**C**) in the hindbrain region of the fish is seen 3 hpi. (**D**) Overlay of (**B**) and (**C**). (**E**–**G**) colocalization is seen between **6a** (**E**) and astrocytes (**F**) in the telencephalon region of the fish. (**G**) Overlay of (**E**) and (**F**). (**H**) Similarly, first generation probe, Rh_B_MP crosses the BBB to label astrocytes in the telencephalon. The control molecule, MitoTracker, is seen in the vasculature, but not in the brain, in both telencephalon (**I**) and dorsal to optic tectum (**J**) region. Arrowheads indicate vasculature labeling outside of the brain. Scale = 25 μm.

In summary, we have synthesized new astrocyte-specific fluorophores and have shown that our small library selectively marks mitochondria and has the ability to traverse the BBB in zebrafish. We anticipate expanding the utility of this targeting method to additional fluorophores that span the visible and infrared-light spectrum so long as they can be modified to achieve a permanent +2 charge containing the methyl-pyridinium astrocyte-targeting moiety. Additionally, these experimental results suggest there is potential to add this targeting moiety to other biologically relevant molecules like transcription activators, drugs, or calcium indicators for targeted delivery to astrocytes, or where an increase in BBB permeability is desired, since delivery of molecules to the brain is a central challenge faced in drug-delivery. Traceless delivery of small molecules that can be uncaged using light or enzyme stimuli is of particular interest, resulting in spatiotemporal control of fluorescent imaging, drug delivery, or transcription activation. It is also of use to researchers to classify sub-populations of astrocytes considering their heterogenous nature, in order to inform the diverse functions of these cells. Elucidating this sub-population and obtaining more information about the mechanisms of transport into astrocytes is of particular interest, since it can help guide further cargo diversification.

## ASSOCIATED CONTENT

### Supporting Information

The Supporting Information is available free of charge via the Internet at http://pubs.acs.org. The Supporting Information includes a list of abbreviations, materials and methods, primary mouse astrocyte cortical culture, HEK293T and HeLa cell culture, zebrafish ventricle injection, zebrafish pericardial injection, general synthetic schemes, Supplementary Figures 1–10, Supplementary Schemes 1–2, Supplemental Movie 1.

## AUTHOR INFORMATION

### Present Addresses

†If an author’s address is different than the one given in the affiliation line, this information may be included here.

### Author Contributions

†D.A.C. and N.G. contributed equally. D.A.C. performed experiments, analyzed data, and wrote the manuscript. N.G. performed experiments, analyzed data, and critically read the manuscript. S.S. and O.H. performed experiments and critically read the manuscript. S.T.L. conceived the project and wrote the manuscript. The manuscript was written through contributions of all authors. / All authors have given approval to the final version of the manuscript.

### Notes

Any additional relevant notes should be placed here.

## ACKNOWLEDGMENT

We thank both the Tsirka and Colognato labs for help with mouse primary cells, Nan Wang for high resolution mass spectrometry analysis, the Stony Brook NMR facility for use of NMR instruments, and the Tonge, Carrico, and Sampson groups.

## ABBREVIATIONS

BBB: blood-brain-barrier
CNS: central nervous system
DMF: dimethylformamide
dpf: days post-fertilization
GFAP: glial fibrillary acidic protein
GFP: green fluorescent protein
hpi: hours-post injection
OCT: Organic Cation Transporter
HATU: 1-[bis(dimethylamino)methylene]-1H-1,2,3-triazolo[4,5-b]pyridinium 3-oxid hexafluorophosphate
MCC: Mander’s colocalization coefficient
mM: millimolar
MTT: 3-(4,5-dimethylthiazol-2-yl)-2,5-diphenyltetrazoliumbromide
PTU: phenylthiourea
μM: micromolar

## REFERENCES

1. Herculano-Houzel, S. The human brain in numbers: a linearly scaled-up primate brain. Front Hum Neurosci 3, 31 (2009).

2. Benarroch, E. E. Neuron-Astrocyte Interactions: Partnership for Normal Function and Disease in the Central Nervous System. Mayo Clin Proc 80, 1326–1338 (2005).

3. Verkhratsky, A., Zorec, R. & Parpura, V. Stratification of astrocytes in healthy and diseased brain. Brain Pathology 27, 629–644 (2017).

4. Sofroniew, M. V. & Vinters, H. V. Astrocytes: biology and pathology. Acta Neuropathol 119, 7–35 (2010).

5. Nolte, C. et al. GFAP promoter-controlled EGFP-expressing transgenic mice: A tool to visualize astrocytes and astrogliosis in living brain tissue. Glia 33, 72–86 (2001).

6. Cahoy, J. D. et al. A Transcriptome Database for Astrocytes, Neurons, and Oligodendrocytes: A New Resource for Understanding Brain Development and Function. The Journal of Neuroscience 28, 264–278 (2008).

7. Ludwin, S. K., Kosek, J. C. & Eng, L. F. The topographical distribution of S-100 and GFA proteins in the adult rat brain: An immunohistochemical study using horseradish peroxidase-labelled antibodies. Journal of Comparative Neurology 165, 197–207 (1976).

8. Zuo, Y. et al. Fluorescent Proteins Expressed in Mouse Transgenic Lines Mark Subsets of Glia, Neurons, Macrophages, and Dendritic Cells for Vital Examination. 24, 10999–11099 (2004).

9. Oberheim, N. A., Goldman, S. A. & Nedergaard, M. Heterogeneity of astrocytic form and function. Methods in Molecular Biology 814, 23–45 (2012).

10. Oberheim, N. A., Wang, X., Goldman, S. & Nedergaard, M. Astrocytic complexity distinguishes the human brain. Trends Neurosci 29, 547–553 (2006).

11. Nimmerjahn, A., Kirchhoff, F., Kerr, J. N. D. & Helmchen, F. Sulforhodamine 101 as a specific marker of astroglia in the neocortex in vivo. Nat Methods 1, 31–37 (2004).

12. Schnell, C., Hagos, Y. & Hülsmann, S. Active sulforhodamine 101 uptake into hippocampal astrocytes. PLoS One 7, e49398 (2012).

13. Zimmermann, M. & Stan, A. C. PepT2 transporter protein expression in human neoplastic glial cells and mediation of fluorescently tagged dipeptide derivative β-Ala-Lys-Nε-7-amino-4-methyl-coumarin-3-acetic acid accumulation. J Neurosurg 112, 1005–1014 (2010).

14. Santos, F. M. F. et al. Cyanine-Like Boronic Acid-Derived Salicylidenehydrazone Complexes (Cy-BASHY) for Bioimaging Applications. Chemistry – A European Journal 26, 14064–14069 (2020).

15. Preston, A. N., Cervasio, D. A. & Laughlin, S. T. Visualizing the brain’s astrocytes. Methods Enzymol 622, 129–151 (2019).

16. Preston, A. N. et al. Visualizing the Brain’s Astrocytes with Diverse Chemical Scaffolds. ACS Chem Biol 13, 1493–1498 (2018).

17. Preston, A. N. et al. Design Principles for Cationic, Astrocyte-Targeted Probes. ChemBioChem 20, 366–370 (2019).

18. Grimm, J. B. et al. Carbofluoresceins and carborhodamines as scaffolds for high-contrast fluorogenic probes. ACS Chem Biol 8, 1303–1310 (2013).

19. Grimm, J. B. et al. A general method to finetune fluorophores for live-cell and in vivo imaging. Nat Methods 14, 987 (2017).

20. Grimm, J. B. et al. A general method to improve fluorophores for live-cell and single-molecule microscopy. Nat Methods 12, 244 (2015).

21. Grimm, J. B. & Lavis, L. D. Synthesis of rhodamines from fluoresceins using pd-catalyzed c-n cross-coupling. Org Lett 13, 6354–6357 (2011).

22. Johnson, L. V., Walsh, M. L. & Chen, L. B. Localization of mitochondria in living cells with rhodamine 123. Proc Natl Acad Sci U S A 77, 990–994 (1980).

23. Dunn, K. W., Kamocka, M. M. & Mcdonald, J. H. A practical guide to evaluating colocalization in biological microscopy. Am J Physiol Cell Physiol 300, 723–742 (2011).

24. Manders, E. M. M., Verbeek, F. J. & Aten, J. A. Measurement of co-localization of objects in dual-colour confocal images. J Microsc 169, 375–382 (1993).

25. Poskanzer, K. E. & Yuste, R. Astrocytes regulate cortical state switching in vivo. Proc Natl Acad Sci U S A 113, E2675–84 (2016).

26. Mu, Y. et al. Glia Accumulate Evidence that Actions Are Futile and Suppress Unsuccessful Behavior. Cell (2019) doi:10.1016/J.CELL.2019.05.050.

27. Chen, J., Poskanzer, K. E., Freeman, M. R. & Monk, K. R. Live-imaging of astrocyte morphogenesis and function in zebrafish neural circuits. Nature Neuroscience 2020 23:10 23, 1297–1306 (2020).

28. Burgess, H. A. & Granato, M. Modulation of locomotor activity in larval zebrafish during light adaptation. Journal of Experimental Biology 210, 2526–2539 (2007).

29. de Esch, C. et al. Locomotor activity assay in zebrafish larvae: Influence of age, strain and ethanol. Neurotoxicol Teratol 34, 425–433 (2012).

30. Fleming, A., Diekmann, H. & Goldsmith, P. Functional Characterisation of the Maturation of the Blood-Brain Barrier in Larval Zebrafish. PLoS One 8, e77548 (2013).

31. Kim, S. S. et al. Zebrafish as a Screening Model for Testing the Permeability of Blood-Brain Barrier to Small Molecules. Zebrafish 14, 322–330 (2017).

32. Quinonez-Silvero, C., Hübner, K. & Herzog, W. Development of the brain vasculature and the blood-brain barrier in zebrafish. Dev Biol 457, 181–190 (2020).

33. Umans, R. A. et al. CNS angiogenesis and bar-riergenesis occur simultaneously. Dev Biol 425, 101–108 (2017).

