## Supplemental Information for "Blood-brain-barrier permeable fluorescent astrocyte probes"

### Table of Contents

**List of abbreviations** ... 3
**General materials and methods** ... 4
**Experiments involving HEK293T and HeLa cells** ... 5
Administration of compounds by bathing ... 5
MTT Assay ... 5
**Experiments involving primary astrocyte culture** ... 6
Administration of compounds by bathing ... 6
Labelling with Mito Tracker ... 6
**Experiments involving zebrafish larvae** ... 7
Zebrafish microinjection ... 7
Light-evoked behavioural assay ... 7
**Supplementary Figure 1** ... 8
**Supplementary Figure 2** ... 9
**Supplementary Figure 3** ... 10
**Supplementary Figure 4** ... 11
**Supplementary Figure 5** ... 12
**Supplementary Figure 6** ... 13
**Supplementary Figure 7** ... 14
**Supplementary Figure 8** ... 15
**Supplementary Figure 9** ... 16
**Supplementary Figure 10** ... 17
**Supplementary Scheme 1: General scheme for synthesis of methylpyridinium linker** ... 18
**Supplementary Scheme 2: General scheme for the synthesis of modified Janelia Fluors®** ... 18
$^{1}$H and $^{13}$C NMR spectra for **4a** ... 20
$^{1}$H and $^{13}$C NMR spectra for **5a** ... 22
$^{1}$H and $^{13}$C NMR spectra for **6a** ... 24
**References** ... 26

**List of abbreviations**

| | |
|---|---|
| BBB | Blood-brain-barrier |
| DCM | Dichloromethane |
| DIPEA | N,N-diisopropylethylamine |
| DMF | Dimethylformamide |
| dpf | Days post fertilization |
| E3 | Embryo media |
| ESI | Electrospray ionization |
| EtOAc | Ethyl acetate |
| EtOH | Ethanol |
| FBS | Fetal bovine serum |
| HATU | Hexafluorophosphate Azabenzotriazole Tetramethyl Uronium |
| Hpf | Hours-post-fertilization |
| Hpi | Hours-post injection |
| HPLC | High pressure liquid chromatography |
| HRMS | High resolution mass spectrometry |
| MHz | Mega Hertz |
| MeCN | Acetonitrile |
| MeOH | Methanol |
| MS | Mass spectrometry |
| ms | Millisecond |
| PTU | Phenylthiourea |
| $R_t$ | Retention time |
| TEA | Triethylamine |
| TFA | Trifluoroacetic acid |
| THF | Tetrahydrofuran |
| TLC | Thin layer chromatography |
| UV | Ultraviolet |

**General materials and methods**.

All chemical reagents were of analytical grade, obtained from commercial suppliers, and used without further purification unless otherwise specified. Reactions were monitored by thin layer chromatography (TLC) on pre-coated glass TLC plates (Analtech UNIPLATE™ silica gel HLF w/ organic binder, 250 µm thickness, with UV254 indicator). TLC plates were visualized by UV illumination. Flash chromatography was performed using Sorbtech, 60 Å, 40–63 µm or Millipore 60 Å, 35–70 µm silica gel according to the procedure described by Still[1]. HPLC was performed using a Shimadzu HPLC (FCV-200AL) equipped with an Agilent reversed phase Zorbax Sb-Aq C18 column (4.6×250 mm or 21.2×250 mm) fitted with an Agilent stand-alone prep guard column. NMR spectra ($^{1}H$ and $^{13}C$) were obtained using a 400, 500, or 700 MHz Bruker spectrometer and analyzed using Mestrenova 9.0. $^{1}H$ and $^{13}C$ chemical shifts (δ) were referenced to residual solvent peaks. Low-resolution electrospray ionization (ESI) mass spectra and high-resolution (ESI) mass spectra were obtained at the Stony Brook University CASDA Mass Spectrometry Center with an Agilent 6110 Single Quad LC/MSD and Bruker Impact II QTOF spectrometer, respectively. All cells were live-imaged in L-15 media using a Zeiss Axio Examiner.D1 modified with an Andor Differential Scanning Disk confocal unit and a 40x NA 1.0 or 20x NA 0.5 water immersion objective. Confocal images were analyzed using ImageJ (NIH) and prepared for presentation using Adobe Illustrator. The same brightness and contrast settings were applied to all images in a given figure, for a given molecule.

The fidelity of all DNA constructs was confirmed by DNA sequencing at the Stony Brook University DNA Sequencing facility. Oligonucleotides were purchased from Integrated DNA Technologies (Coralville, IA). TOP10 One-Shot and Mach1 competent cells, Lipofectamine 3000, and cell culture media were purchased from Invitrogen (Carlsbad, CA). Millicell EZ 8-chamber slides were purchased from Millipore EMD (Darmstadt, Germany). Phusion DNA polymerase, Taq DNA polymerase, calf intestinal alkaline phosphatase (CIP), and restriction enzymes were purchased from New England BioLabs (Beverly, MA). Omega E.Z.N.A.® gel extraction kit and Omega E.Z.N.A.® plasmid mini kit II were purchased from Omega Bio-Tek, Inc. (Norcross, GA).

#### Experiments involving HEK293T and HeLa cells in culture

##### Culturing of HEK293T and HeLa cells

HEK293T or HeLa cells were maintained in high-glucose Dulbecco's Modified Eagle Medium (DMEM) supplemented with 10% FBS and 1% antibiotic-antimycotic solution at 37 °C in air with 5% $CO_2$. Cells were passaged on alternating days when they reached approximately 90% confluency. Cells were passaged into poly-d-lysine-coated Fisherbrand 6-well plates at 300,000 cells/well, cultured for 24 h then transfected with EGFP control or OCT1-p2a-EGFP using Lipofectamine 3000 according to the manufacturer's instructions.

##### Administration of compounds by bathing

After the removal of transfection media, 6-well plates were bathed in warm DMEM with 1 μM test molecule. The cells were incubated at 37 °C in air with 5% $CO_2$ for 20 min after which bathing solutions were removed and 1 mL of warm HBSS was added to each well. The HBSS was removed, replaced with 1 μg/mL Hoescht 33342 in L-15 media and incubated at 37 °C in air with 5% $CO_2$. Cells were imaged after 30-45 mins using a Zeiss Axio Examiner.D1 modified with an Andor Differential Scanning Disk confocal unit and a 40x NA 1.0 water immersion objective. Parameters adjusted to optimize image quality are the exposure time of the camera and the z-range to take a stack of the entire cell.

##### Image analysis

Confocal images were separated into individual channels and a maximum intensity projection of the appropriate channel was made. Brightness and contrast settings for each channel were adjusted consistently across images of the same experiment using ImageJ.

##### MTT Assay

The toxicity of all compounds was evaluated in HEK293T cells. Cells were passaged at approximately 85% confluency into poly-d-lysine-coated 10 cm plates, cultured for 24 h then transfected with EGFP control or OCT1-p2a-EGFP using Lipofectamine 3000 according to the manufacturer's instructions. Cells were cultured for 24 h before passaging approximately 25,000 cells onto poly-d-lysine-coated CytoOne 96-well plates. Cells were cultured for 18-24 h at 37 °C in air with 5% $CO_2$ to allow for adherence to the well surface. Next, the culture medium was removed, and 100 μL fresh medium containing each compound at concentrations of 0, 0.01, 0.1, 1.0 and 10 μM was added to the wells. The plates were incubated for 20 min. After 20 min, the supernatant was removed and 90 μL of fresh DMEM was added. The plates were incubated for another 1 hr. Then 10 μL of 5 mg/mL 3-(4,5-dimethylthiazol-2-yl)-2,5-diphenyltetrazoliumbromide (MTT) was added to each well and was incubated for an additional 4 h. Afterwards 100 μL of the solubilization buffer was added and the cells were incubated overnight to dissolve the formazan crystals formed inside the cells. The next day, the absorbance was measured at 570 nm on a BioTek Synergy 2 Plate Reader and the percent viability was calculated. One-way ANOVA in Excel followed by Tukey's HSD at $p = 0.05$ was performed on $n = 3$ trials each containing 6 technical replicates.

#### Experiments involving cultured primary mouse cortical astrocytes

##### Culturing of P2-P8 primary mouse cortical astrocytes

Primary cultures of astrocytes were prepared from P8 C57/Bl6 mice or neonatal p0–2 MacGreen mice, which express GFP under the microglia/macrophage promoter CSF1R, using the protocol previously described by Tsirka and co-workers[2]. Briefly, after chemical dissociation with papain and several resuspension steps, plates were placed on an orbital shaker at days 7–10 to remove microglia and oligodendrocytes. To isolate the astrocytes after separation of microglia and OPCs from the mixed cortical culture, the remaining astrocyte monolayer in the culture was trypsinized and the media was collected with detached astrocytes and maintained in complete media (high glucose DMEM with 10% FBS, and 1% antibiotic-antimycotic solution) on PDL-coated (50 μg/mL) substrates at 37 °C in air with 5% $CO_2$.

##### Culturing of E18 primary mouse cortical astrocytes

Primary cultures were prepared from prenatal C57 E18 cortex obtained through Transnetyx. Cortices were digested using the protocol previously described by Tsirka and co-workers[2]. Briefly, after chemical dissociation with papain and several resuspension steps, plates were placed on an orbital shaker at days 7–10 to remove microglia and oligodendrocytes. To isolate the astrocytes after separation of microglia and OPCs from the mixed cortical culture, the remaining astrocyte monolayer in the culture was trypsinized and the media was collected with detached astrocytes and maintained in complete media (high glucose DMEM with 10% FBS, and 1% antibiotic-antimycotic solution) on PDL-coated (50 μg/mL) substrates at 37 °C in air with 5% $CO_2$.

##### Administration of compounds by bathing

Astrocytes were passaged into a poly-d-lysine coated (50 μg/mL) Fisherbrand 6-well plate or 8-well chamber and seeded at 300,000 cells/well in 2 mL complete media or 15,000 cells/well with 0.4 mL complete media, respectively. A bathing solution of 1 μM test molecule was prepared in fresh complete media. Media was removed from each well and bathing solutions were added. The cells were incubated at 37 °C with 5% $CO_2$ for 20 mins after which bathing solutions were removed and 1 mL of HBSS was added to each well for washing. The HBSS was then removed, replaced with 1 μg/mL Hoescht 33342 in L-15 media and incubated at 37 °C in air with 5% $CO_2$. Cells were live-imaged in L-15 media using a Zeiss Axio Examiner.D1 modified with an Andor Differential Scanning Disk confocal unit and a 40x NA 1.0 water immersion objective after 30-45 mins. Parameters adjusted to optimize image quality are the exposure time of the camera and the z-range to take a stack of the entire cell.

##### Organelle labeling with MitoTracker

Following bathing and wash with HBSS, each tracker was added to complete media with 1 μg/mL Hoescht 33342 and incubated at 37 °C in air with 5% $CO_2$ for 45 minutes. The final concentrations for bathing in complete media are as follows: 200 nM MitoGreen, 400 nM MitoRed. Bathing solutions were then removed, and cells were live imaged in L15 media using a Zeiss Axio Examiner.D1 modified with an Andor Differential Scanning Disk confocal unit and a 40x NA 1.0 objective. Parameters adjusted to optimize image quality are the exposure time of the camera and the z-range to take a stack of the entire cell.

##### Image analysis: quantification of colocalization

Confocal images were separated into individual channels and a maximum intensity projection of the appropriate channel was made. Brightness and contrast were adjusted for each image consistently across experimental conditions that use the same probe (i.e. all 549MP bathing are adjusted to the same brightness/contrast). Brightness/contrast settings may differ between figures. Using the Colocalization_Finder plugin in ImageJ, a Red (fluorophore) and Green (MitoTracker) channel were assigned. The pixel threshold was manually estimated using the 512x512 scatterplot in conjunction with the composite image overlay to minimize background and are reported in the figure legend. Mander's correlation coefficients M1 and M2 were obtained after the threshold was established.

#### Experiments involving zebrafish larvae

##### Zebrafish husbandry

Adult zebrafish were housed at 28.5 ºC on a 14-h light 10-h dark light cycle. Embryos were produced from natural crosses between one male and one female zebrafish (zebrafish lines: wild type strain AB, HuC:GFP, GFAP:GFP, mpeg1:EGFP, Olig2:EGFP). Fertilized embryos are sorted and seeded at 50 embryos per 10 cm petri dish in 1X embryo media (E3) and placed in an incubator (28.5 ºC, 14-h light, 10-h dark light cycle) to mature. At or before 22-hours postfertilization (hpf), embryos were transferred to a solution containing 197 μM phenylthiourea (PTU) in 1X E3. Larval zebrafish were observed for manipulation and microinjection using a Leica MZ 10 F fluorescence dissecting microscope equipped with a 1x PlanApo objective.

##### Intracerebroventricular injections of larval zebrafish

Larval zebrafish at 3-5 dpf were anesthetized using 0.064 M tricaine. Fish were mounted dorsal up in 50 μl of 1.5% low melting point agarose in E3 solution on a microscope slide. Using a Nanoject II auto-nanoliter injector positioned on a micromanipulator, 2.3 nL of 100 μM (**4a**, 549MP and $Rh_B$MP) or 1 mM (**5a**, 503MP and **6a**, 585MP) was injected into the ventricle near the optic tectum of the fish. For mitochondrial colocalization experiments, a solution of 600 μM probe + 400 μM MitoTracker was used for injection (2.3 nL). After injection, E3 media was added on top of the fish. Agarose was removed from around the fish using a scalpel until it was free to swim. Fish were then transferred to a fresh PTU-containing E3 solution to recover for 3 hours before imaging. Zebrafish reporter lines include cell-specific promoters that drive the expression of GFP. These lines include GFAP-GFP (astrocytes), HuC-GFP (neurons), mpeg1-GFP (microglia) and olig2-GFP (oligodendrocytes).

##### Intravenous pericardial vein injections of larval zebrafish

Larval zebrafish at 7 dpf were anesthetized using 0.064 M tricaine. Fish were mounted dorsal left in 50 μl of 1.5% low melting point agarose in E3 solution on a microscope slide. Using a Nanoject II auto-nanoliter injector positioned on a micromanipulator, 2.3 nL of 1 mM (all compounds) was injected into the pericardial vein or pericardial space; the needle was manipulated and the fish was penetrated at the intersection where the pericardium meets the ventro-caudal pharyngeal arches and dorso-rostral yolk sac. Fish were then transferred to a fresh PTU-containing E3 solution to recover for 3 hours before imaging.

##### Microscopic imaging of larval zebrafish

Larval zebrafish were viewed using a fluorescence dissecting microscope during injection to determine if loading occurred. To image using confocal microscopy, fish were mounted dorsal in 50 μl of a 2% low melting point agarose in E3 solution on a microscope slide. Fish were live-imaged using a Zeiss Axio Examiner.D1 modified with an Andor Differential Scanning Disk confocal unit and a 40x NA 1.0 or 20x NA 0.5 water-immersion objective. Parameters adjusted to optimize image quality are the exposure time of the camera and the z-range to take a stack of the desired region of each fish.

##### Zebrafish confocal image analysis

Confocal images were separated into individual channels using ImageJ software. A maximum Z-projection of 2-5 stacks was made, and brightness/contrast settings were adjusted consistently across experiments, differing only between fluorophores and transgenic fish lines.

##### Zebrafish spontaneous and light-evoked behavioral swim assays

Fish were mounted in 1.5% low melting point agarose and injected with 2.3 nL of a 100 μM ($Rh_B$MP, n = 10), 549MP, n = 10), 1 mM (585MP, n = 10) solution, or a water control (n = 10) at 5 dpf. Behaviors of 6 dpf larvae were recorded the next day using a Zebrabox imaging system (Viewpoint Life Sciences, France) with constant illumination by infrared light and tracking with automated video-tracking software (Zebralab; Viewpoint Life Sciences, France). All experiments were conducted between 12 and 6 pm The visual-motor behavior paradigm consisted of 20 min of acclimation, 15 min in the light followed by a stimulus (light change), and 15 min in the dark. We tracked behavioral parameters such as number, distance, duration, of movements as well as swim time, swim length, and swim speed. Data were assessed in 1-min bins for the analysis of spontaneous movement. One-way ANOVA in Excel followed by Tukey's HSD test at p = 0.01 was performed to assess the significance between treatment conditions (n = 10) during acclimation, light, and dark periods.

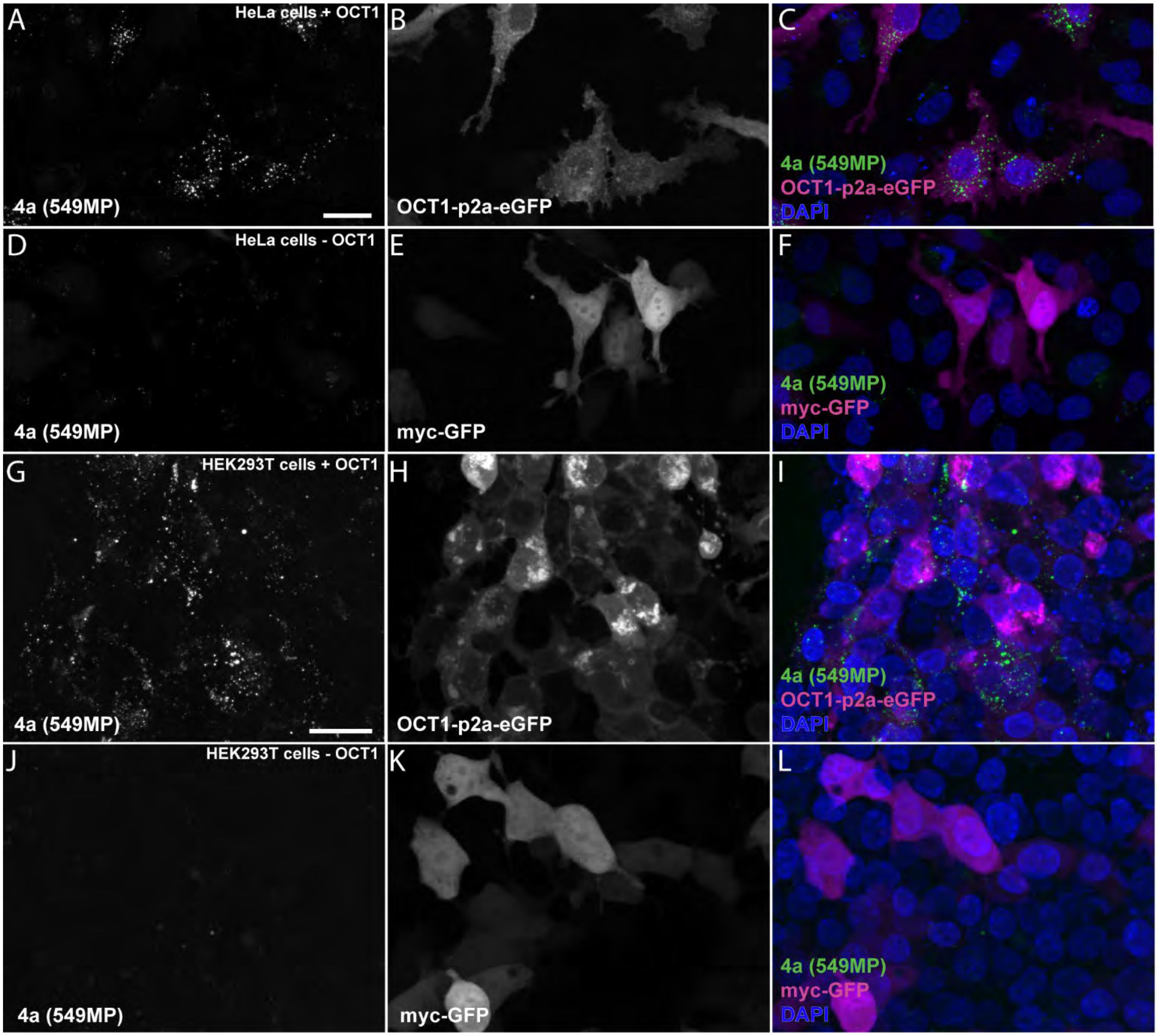


**Supplementary Figure 1. Labeling of HEK293T and HeLa cells expressing OCT1 with 549MP.** After 20 mins incubation with ***4a*** (549MP), only HeLa cells expressing the OCT1 construct take up the molecule (**A-C**), while cells expressing myc-GFP control plasmid cannot (**D-F**). HEK293T cells expressing OCT1 (**G-I**) plasmid take up ***4a***, while HEK293T cells expressing control myc-GFP construct (**J-L**) are unable after 20 mins bathing. Scale = 25 μm.

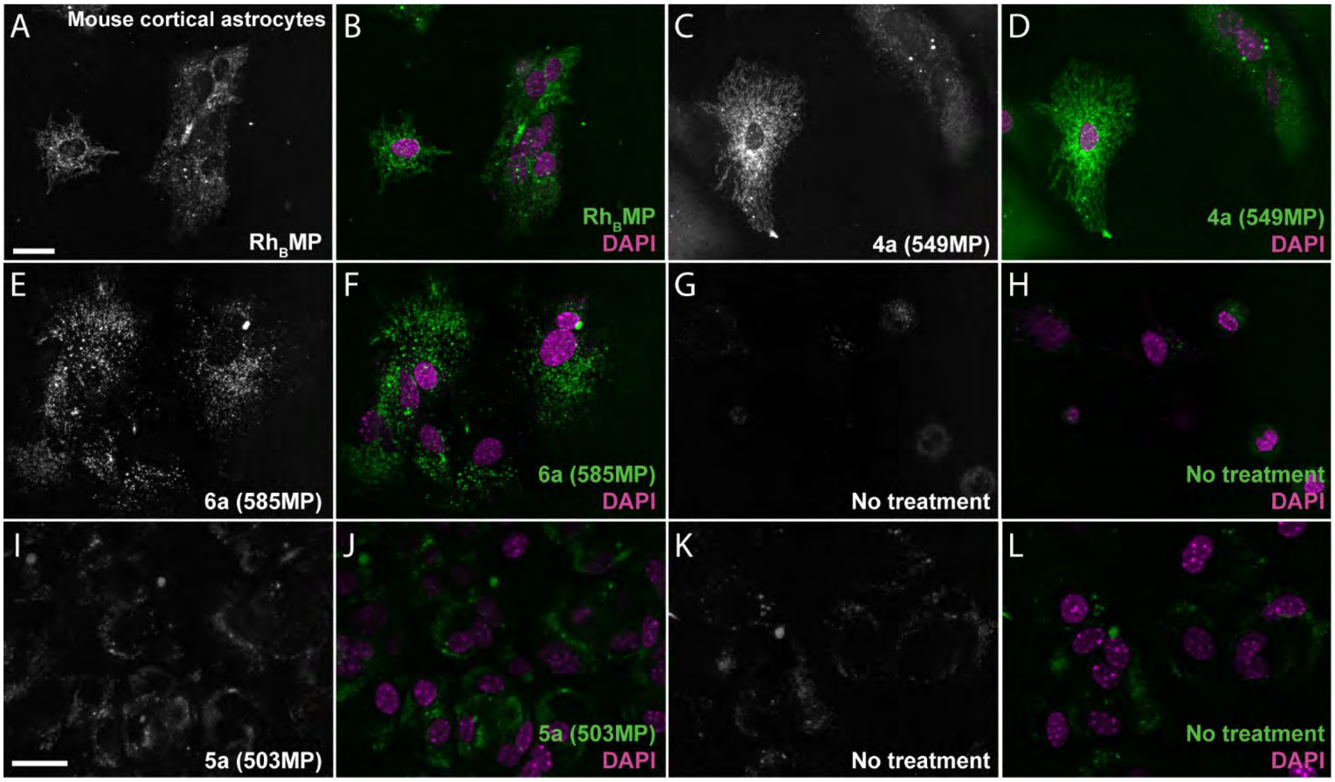


**Supplementary Figure 2. Labeling of mouse cortical astrocytes with 549MP, 585MP, 503MP, and $Rh_B$MP probes.** Mouse cortical astrocytes are labeled with $Rh_B$MP (**A-B**); ***4a***, 549MP (**C-D**); and ***6a***, 585MP (**E-F**) in comparison to autofluorescence seen in the no treatment control (**G-H**) for the red channel after 20 mins incubation at 1 μM. Astrocytes bathed with the green-shifted ***5a***, 503MP at the same concentration (**I-J**) do not seem to show labeling over the green-channel control (**K-L**). Scale = 25 μm.

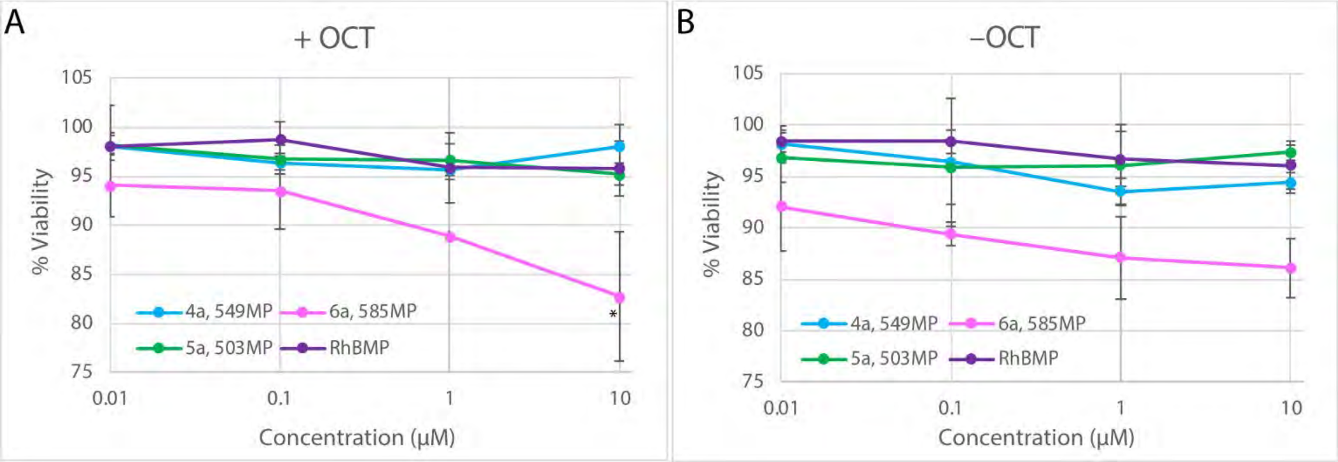


**Supplemental Figure 3. No toxicity is seen in HEK293T cells at normal bathing concentrations of fluorescent probes.** Percent viability after 20 mins bathing for +OCT (**A**) and –OCT (**B**) conditions. Significance is seen only with compound ***6a*** between 0.01 µM and 10 µM conditions in **A**. One way ANOVA, Tukey's HSD, *p = 0.05, n = 3.

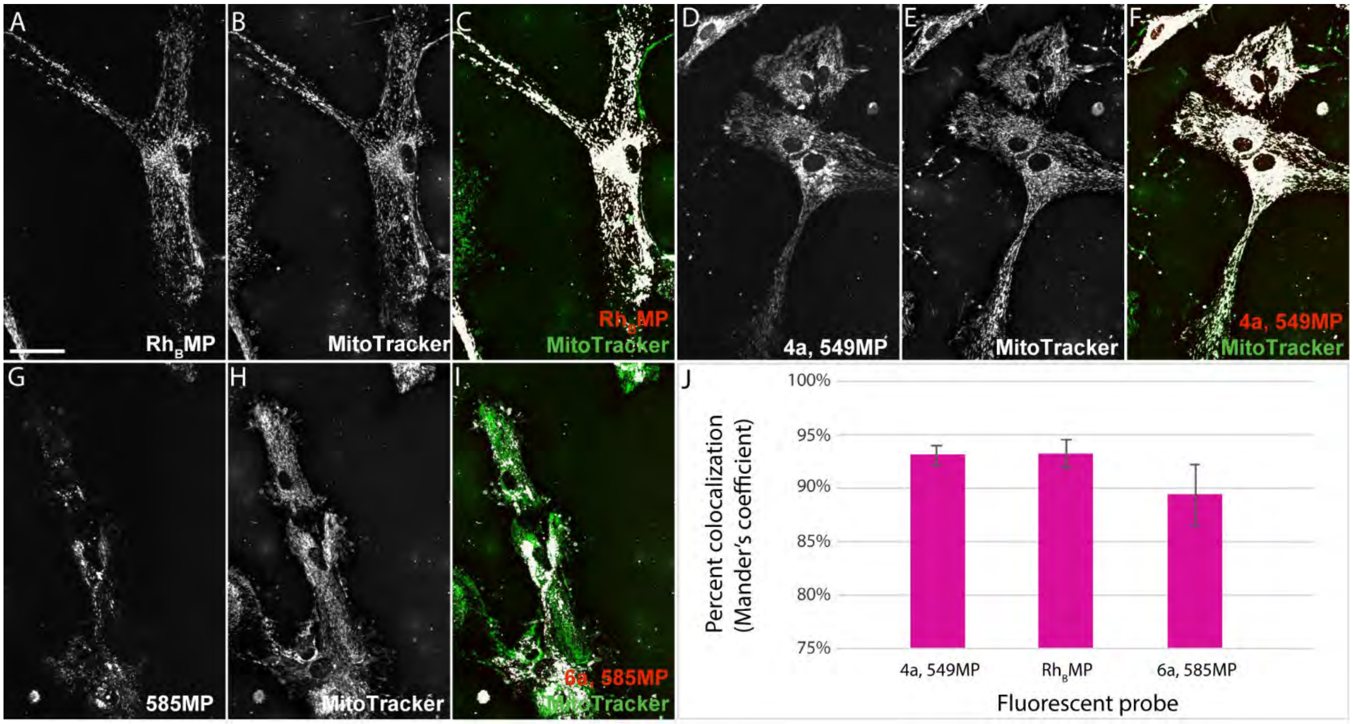


**Supplementary Figure 4. Mitochondrial colocalization between $Rh_B$MP, *4a* (549MP), and *6a* (585MP).** Clear colocalization is seen between $Rh_B$MP (**A**), and MitoTracker (**B**) as shown in (**C**) via the white color which indicates overlapping pixels (pixel threshold x, 282; y, 399). Mander's coefficient for $Rh_B$MP: M1 = 0.928, M2 = 0.661. Colocalization is seen between ***4a,*** 549MP (**D**) and MitoTracker (**E**) as shown via the white overlapping pixels in (**F**) (pixel threshold x, 195; y, 529). Mander's coefficient for ***4a***: M1 = 0.904, M2 = 0.785. Colocalization, albeit less robust, is also seen between ***6a,*** 585MP (**G**) and MitoGreen (**H**) shown by the overlapping pixels in (**I**) (pixel threshold x, 129; y, 295). Mander's coefficient for ***6a***: M1 = 0.950, M2 = 0.259. (**J**) Percent colocalization of fluorophore and MitoTracker for n = 3 replicates. Mander's colocalization coefficient (MCC) is a measurement of colocalization that gives the percentage of pixels in image A (red channel, all probes) that overlap with image B (green channel, MitoTracker), called M1, or the percentage of pixels in image B that overlap with image A, called M2. All MCC values were derived from the "Colocalization_Finder" plugin for imageJ[3]. Pixel threshold in x,y plane was calculated manually to minimize background by selecting pixels above zero in the correlation diagram which created panels **C**, **F**, and **I** as output for $Rh_B$MP, ***4a***, and ***6a***, respectively. Scale bar = 25 µm.

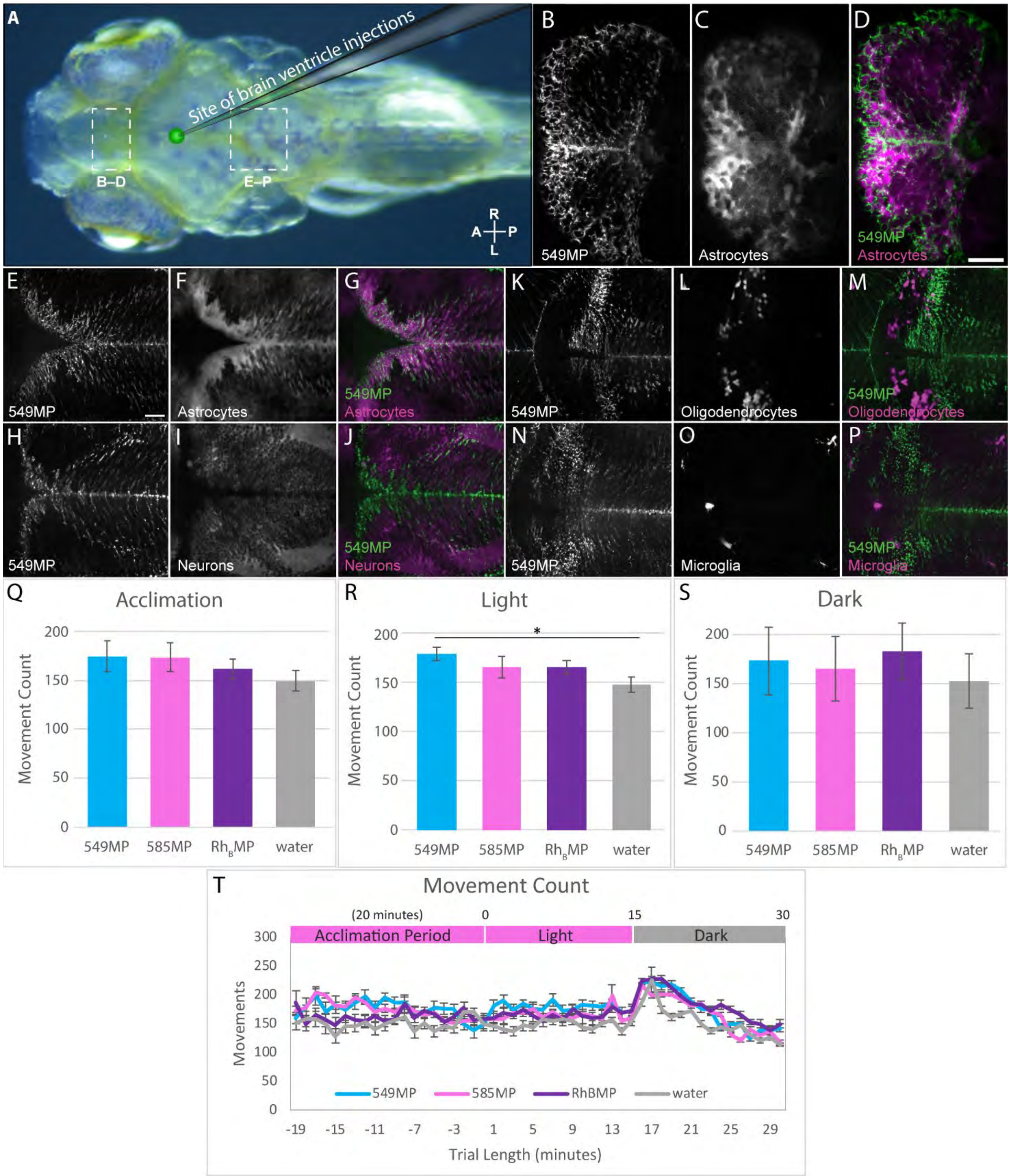


**Supplementary Figure 5. 549MP labels astrocytes in the telencephalon and hindbrain, but not other glial cells or neurons following ventricle injection in larval zebrafish.** After direct ventricle injection (**A**) of ***4a***, 549MP (2.3 nL of a 100 μM solution) into 3-5 dpf zebrafish brain, colocalization is seen with astrocytes in the telencephalon (**B-D**) and hindbrain (**E-G**). Colocalization is not seen between ***4a*** and oligodendrocytes (**K-M**), neurons (**H-J**), or microglia (**N-P**). Zebrafish driver lines are GFAP-GFP (astrocytes), olig2-GFP (oligodendrocytes), mpeg1-GFP (microglia), and HuC-GFP (neurons) which are pseudo-colored magenta in these images. Average movement count in the acclimation (**Q**), light (**R**), and dark (**S**) period as measured via a 50 min behavioral assay of 6 dpf larvae at 18 hpi as represented in line graph, **T**. One-way ANOVA, Tukey's HSD, *p = 0.01, n= 10 fish per condition. Scale = 25 μm.

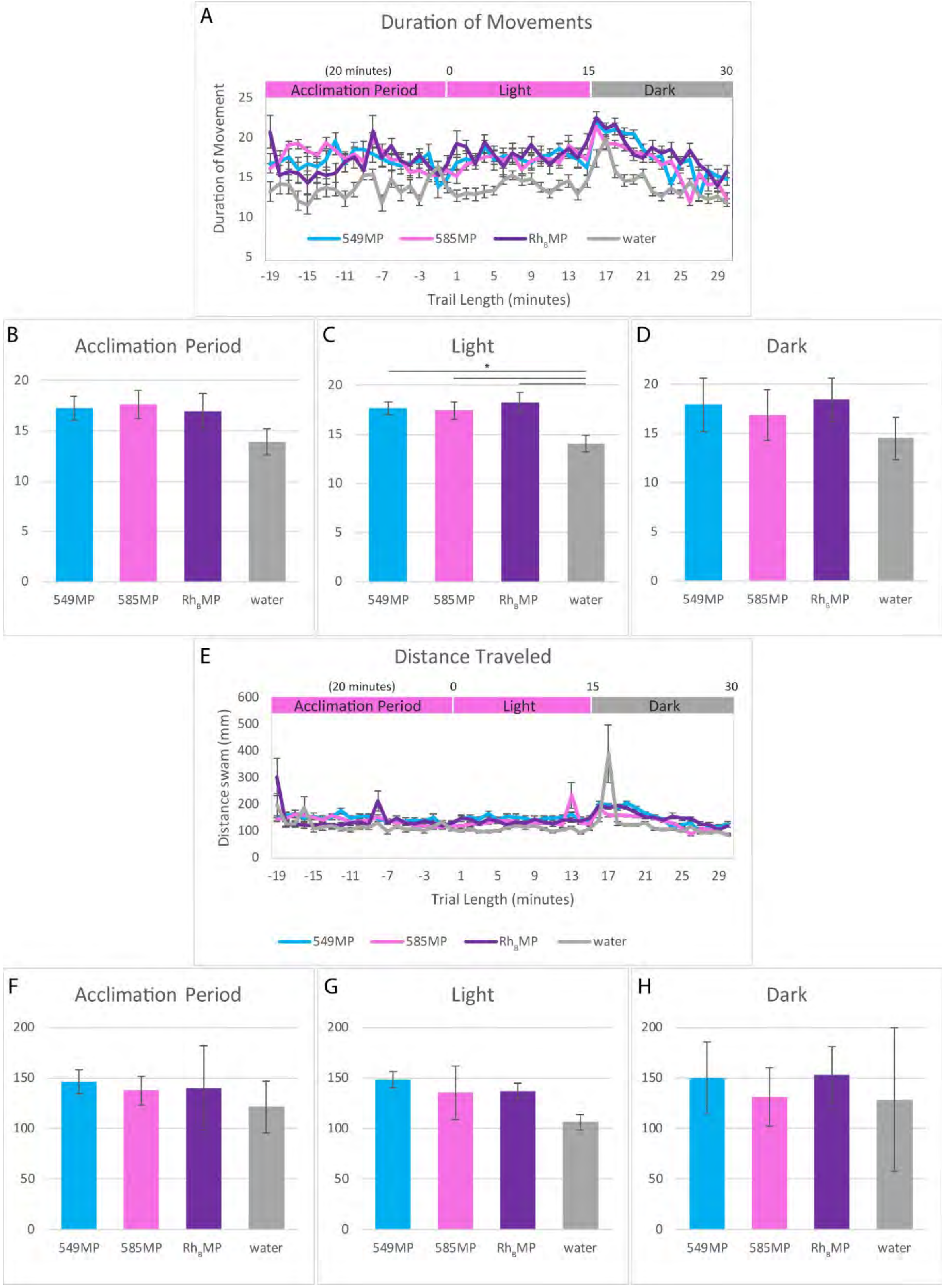


**Supplementary Figure 6. Larval zebrafish at 6 dpf injected with fluorescent probes do not display deficits in movement parameters**. Duration of movement (**A-D**) and distance traveled (**E-H**) was measured via a spontaneous light-evoked behavioral assay at 18 hours following direct ventricle injection of each probe (2.3 nL of a 100 M solution, ***4a*** 549MP & Rh$_B$MP; or 2.3 nL of a 1 mM solution, ***6a*** 585MP). Duration of movements (secs) throughout the 50 min assay represented as a line graph (**A**) which is broken down into acclimation (**B**), light (**C**), and dark (**D**) periods. A significant difference is seen in the duration of movements only in the 15 min light period between each probe and the water control. Distance traveled in millimeters is represented in line graph (**E**) and broken down further into acclimation (**F**), light (**G**), and dark (**H**) periods where no significant difference is seen. One-way ANOVA, Tukey's HSD, *p = 0.01.

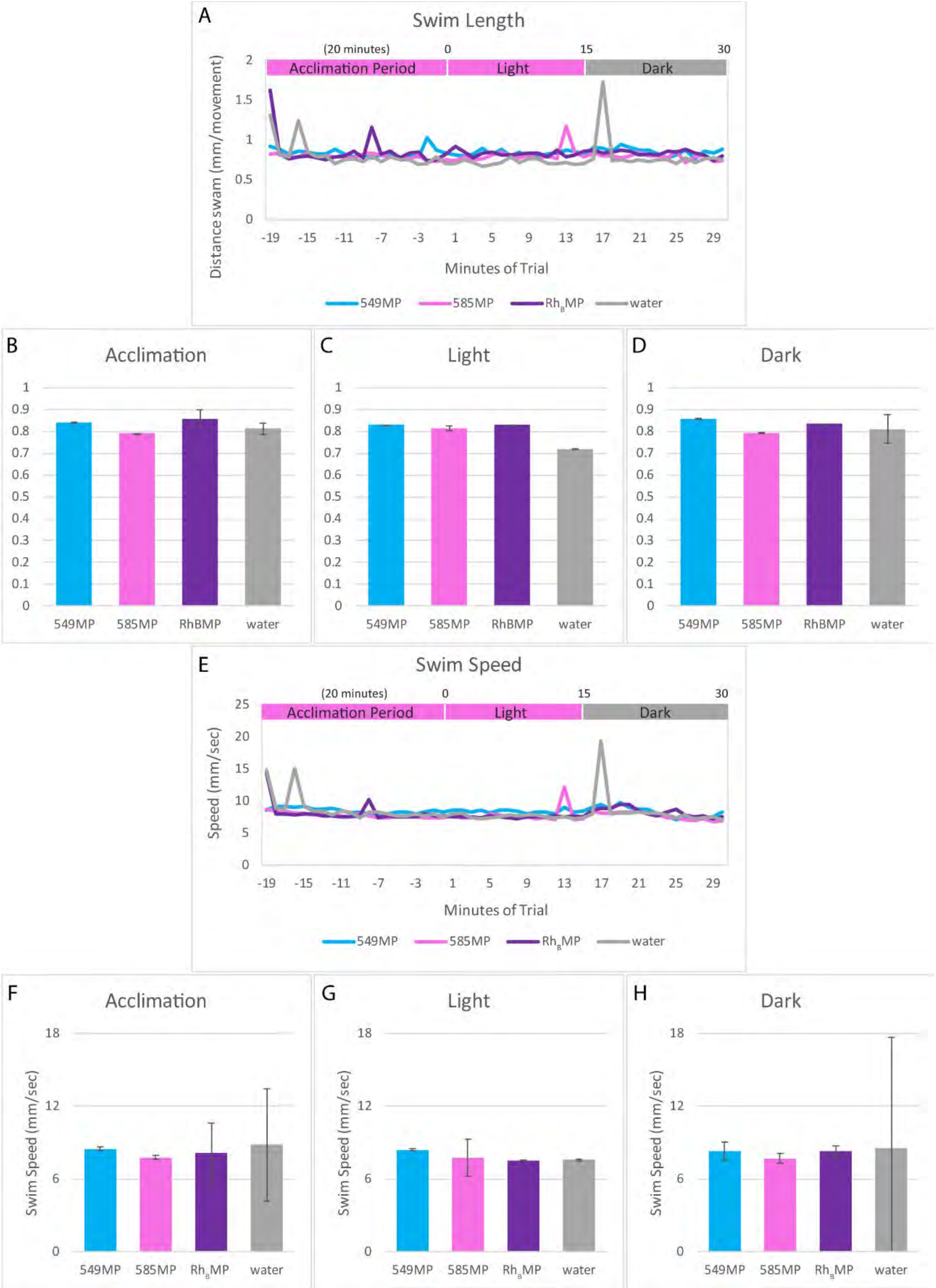


**Supplementary Figure 7. Larval zebrafish at 6 dpf injected with fluorescent probes do not display deficits in movement parameters.** Swim length (**A-D**) and swim speed (**E-H**) were calculated from measured parameters in a spontaneous light-evoked behavioral assay at 18 hours following direct ventricle injection of each probe (2.3 nL of a 100 M solution, ***4a*** 549MP & $Rh_B$MP; or 2.3 nL of a 1 mM solution, ***6a*** 585MP). Swim length is calculated by dividing distance in millimeters by movement count, while swim speed is calculated by dividing distance in millimeters by duration in seconds. Swim length during the 50 min assay is represented as a line graph (**A**) which is broken down into acclimation (**B**), light (**C**), and dark (**D**) periods. Swim speed is represented in line graph (**E**) and broken down further into acclimation (**F**), light (**G**), and dark (**H**) periods. No significant difference is seen. One-way ANOVA, Tukey's HSD, *p = 0.01.

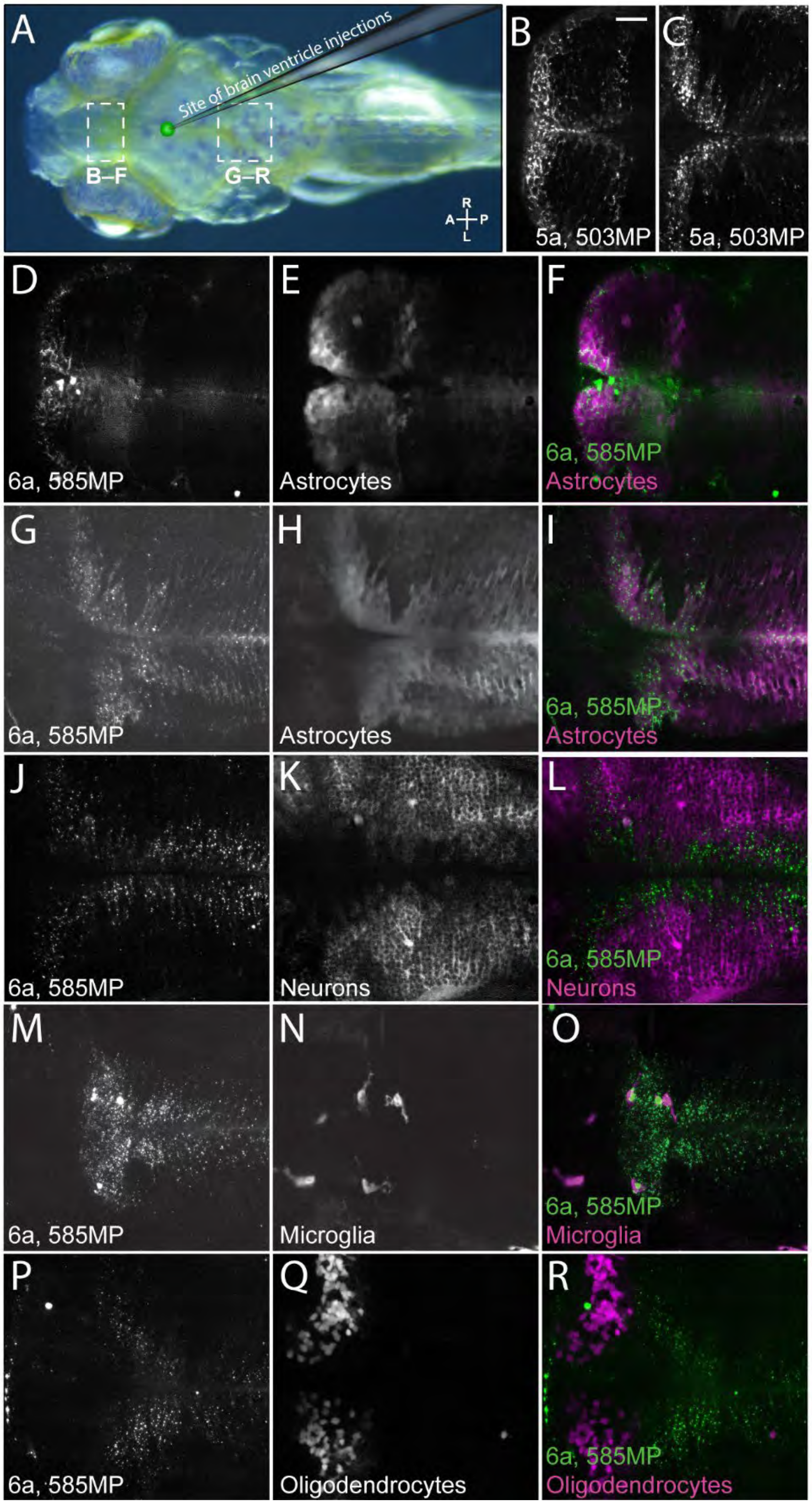


**Supplementary Figure 8. Direct ventricle injection of probes 5a, 503MP and 6a, 585MP reveals a pattern of astrocyte uptake throughout the zebrafish brain.** Direct ventricle injection of ***5a***, 503MP (**B-C**) (2.3 nL of a 1 mM solution) shows a similar pattern as ***6a***, 585MP. Colocalization with zebrafish GFP reporter lines is not possible since ***5a*** is a green-shifted fluorophore. On the other hand, ***6a*** (2.3 nl of a 1 mM solution) shows colocalization between astrocytes in the telencephalon (**D-F**) and hindbrain (**G-I**) regions, but not with neurons (**J-L**), microglia (**M-O**), or oligodendrocytes (**P-R**). It may be of interest to note that microglia seem to phagocytose ***6a*** more readily than other probes, shown via the green puncta seen being engulfed by magenta cells (**O**). Scale = 25 μm.

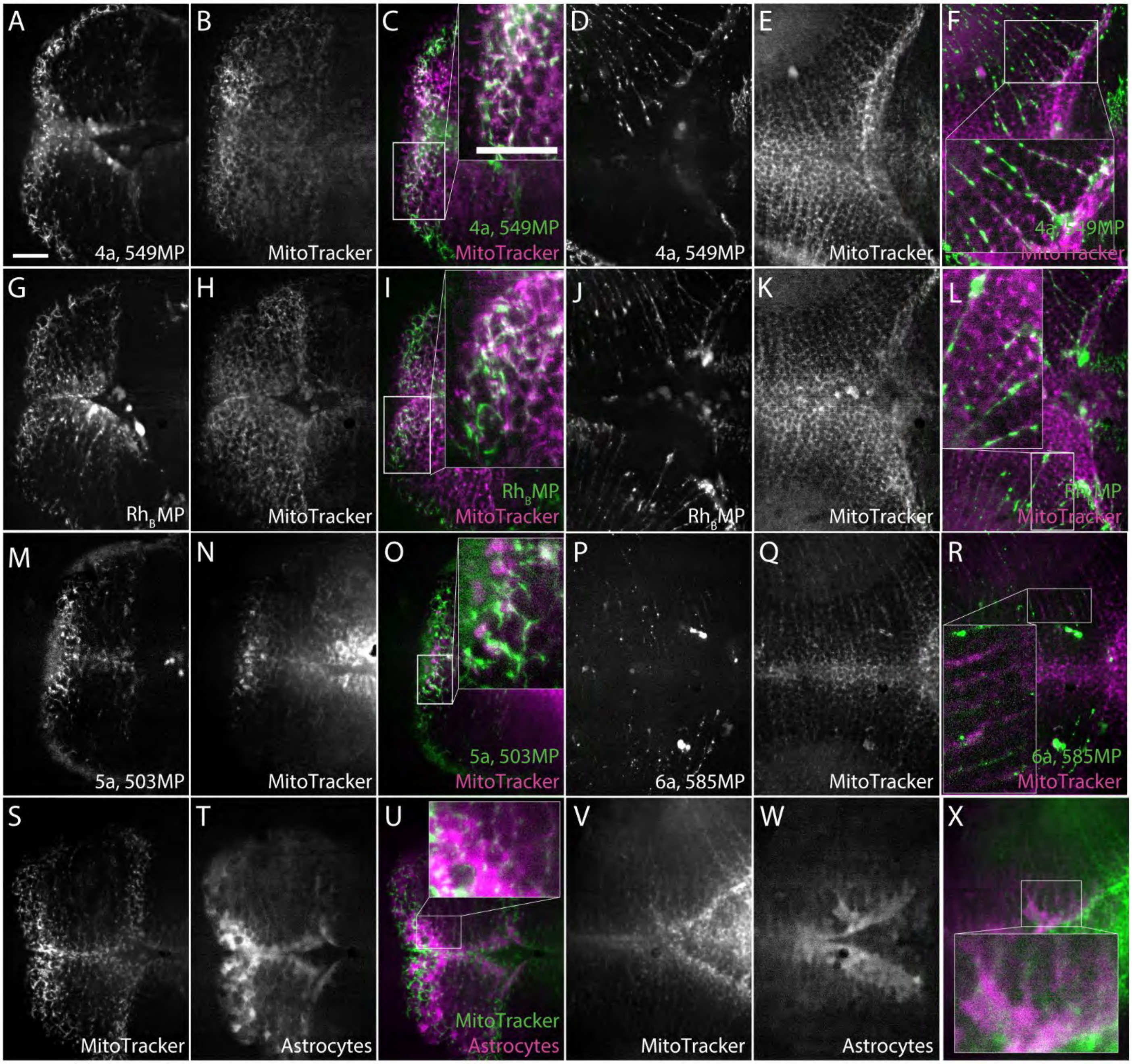


**Supplementary Figure 9. Co-injection of small molecule probes with MitoTracker® into the ventricle reveals colocalization with zebrafish mitochondria**. Direct ventricle injections of a 1 mM solution containing 600 uM ***4a***, 549MP and 400 uM MitoTracker reveal colocalization between zebrafish mitochondria and ***4a*** in the telencephalon (**A-C**) and dorsal to optic tectum (**D-F**). Direct ventricle co-injection of our first-generation probe, $Rh_B$MP (600 uM) and MitoTracker (400 uM) shows colocalization in both telencephalon (**G-I**) and dorsal to optic tectum (**J-L**) of the fish brain. Moreover, ***5a***, 503MP (600 uM) shows a similar pattern in the telencephalon (**M-O**), and ***6a***, 585MP (600 uM) also colocalizes with mitochondria in the optic tectum (**P-R**). Panels **S-X** show MitoTracker labeling of GFAP+ astrocytes in the zebrafish telencephalon (**S-U**) and optic tectum (**V-X**), confirming that our probes label the mitochondria of zebrafish astrocytes. Scale bar = 25 μm.

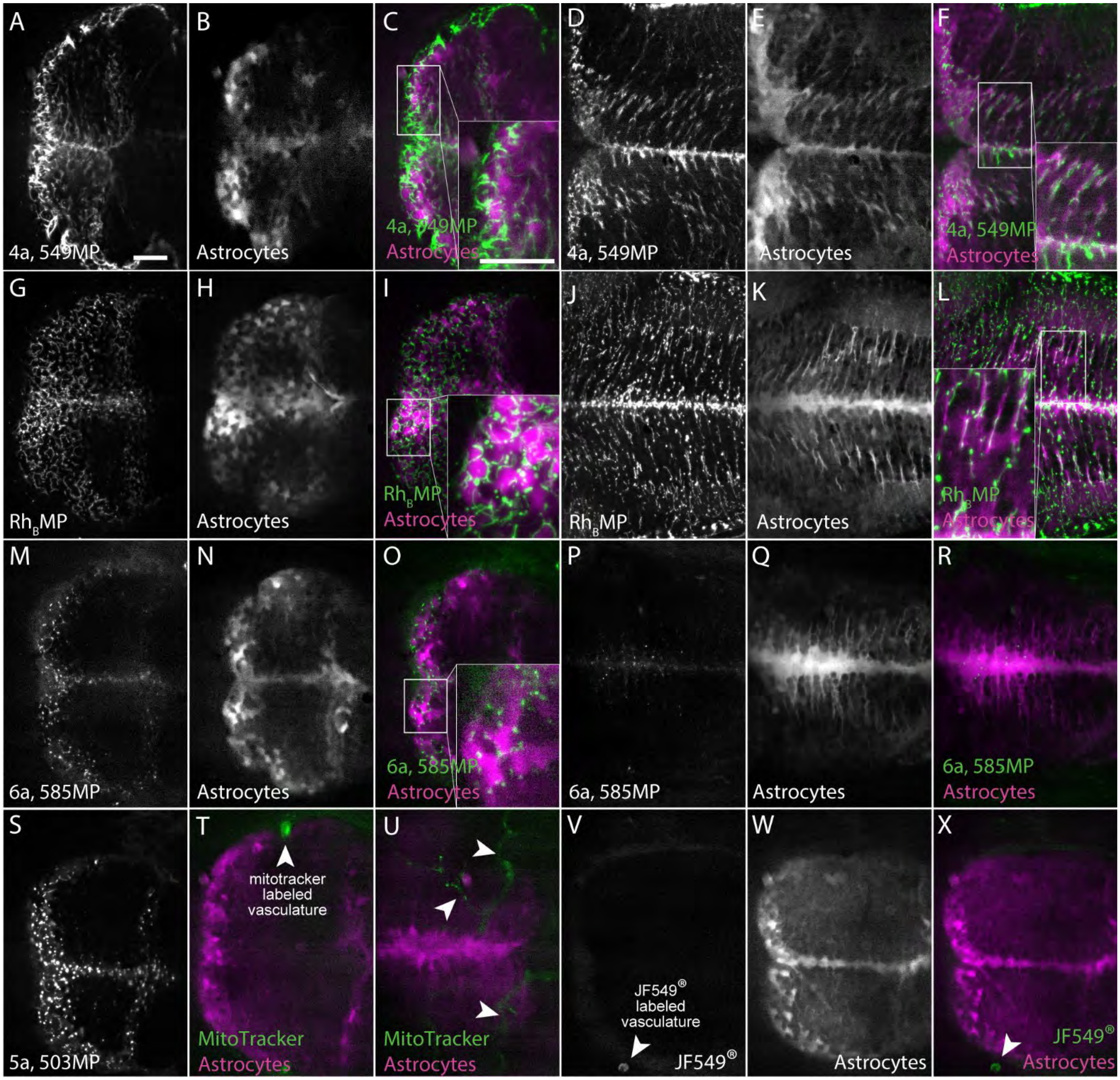


**Supplementary Figure 10. All probes are able to cross the blood-brain-barrier (BBB) after systemic delivery via the pericardial vein in order to reach zebrafish astrocytes.** After pericardial injection (2.3 nL of 1 mM solution) of ***4a***, 549MP into 7 dpf transgenic GFAP-GFP zebrafish (magenta), strong labeling of astrocytes is seen 3 hours-post-injection (hpi) in the telencephalon (**A-C**) and hindbrain (**D-F**) regions. Following systemic introduction of $Rh_B$MP, a similar pattern of astrocyte labeling is seen in both the telencephalon (**G-I**) and hindbrain regions (**J-L**). ***6a***, 585MP is also able to cross the BBB to label astrocytes in the telencephalon (**M-O**) and hindbrain (**P-R**), albeit less robust. ***5a***, 503MP is seen in the telencephalon after systemic injection into WT fish (**S**). A structurally similar control molecule, MitoTracker (**T-U**) is seen retained in the vasculature adjacent to telencephalon (**T**) and dorsal to optic tectum (**U**) after 3 hpi, indicating that the zebrafish BBB is indeed intact for similarly sized molecules. Furthermore, after systemic injection of the unconjugated JF549® (**V-X**), labeling of adjacent vasculature is seen without labeling insinde the telencephalon.  Scale bar = 25 μm.

***Synthesis***

JaneliaFluor® 549 and 503 were synthesized from commercially available fluorescein according to Lavis and coworkers.[4,5] JaneliaFluor® 585 Carbofluorescein was synthesized according to Lavis and coworkers[5,6].

**Supplementary Scheme 1**

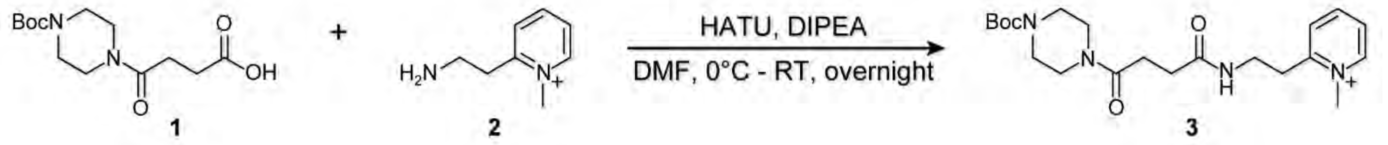


*Synthesis for compounds 1, 2, and 3 have been previously reported*[7,8]. The synthesis of **3** is performed here with some modifications. Briefly, 4-(4-(tert-butoxycarbonyl)piperazin-1-yl)-4-oxobutanoic acid, **1** (1.0 equiv), is dissolved in DMF, cooled to 0°C and both DIPEA (1.2 eq) and HATU (1.2 eq) are added. The mixture is stirred for 30 mins at 0°C. 2-(2-aminoethyl)-1-methylpyridin-1-ium, **2** (1.2 equiv), is added by volume in DMF, and the reaction is stirred overnight at RT. The reaction was concentrated *in vacuo* and purified using previously reported[7] HPLC methods. The Boc protected linker, **3**, is subject to deprotection via 1:1 TFA/DCM and used without further purification in the subsequent coupling reactions between compounds **4-6**.

**Supplementary Scheme 2**

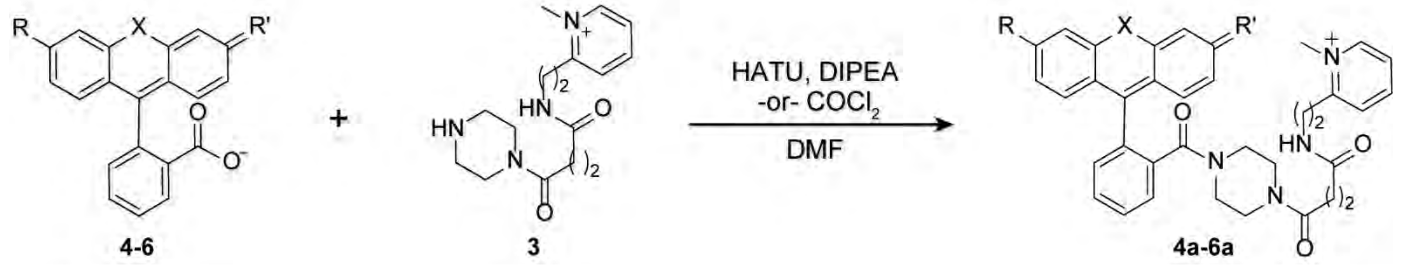


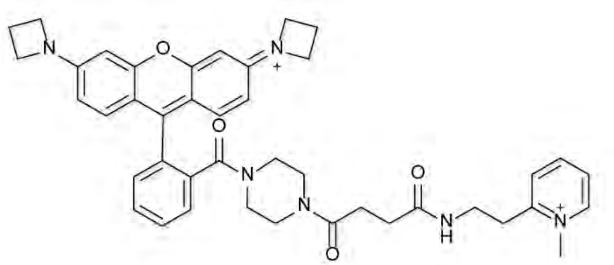

***4a, 549MP:*** JF549 (7.8 mg, 0.019 mmol, 1.0 equiv) was added to a flame-dried flask in 600 µL DCM. DIPEA (12 µL, 0.069 mmol, 3.6 equiv) was added.The reaction was cooled to 0°C and HATU (8.7 mg, 0.023 mmol, 1.2 eq) was added in 450 µL dry DMF. The reaction mixture was stirred at 0°C for 15 minutes. Deprotected piperazine methylpyridinium linker, **3** (5.7 mg, 0.019 mmol, 1 equiv) was dissolved in 300 µL dry DMF, added to the reaction mixture and stirred overnight at RT. The reaction was concentrated in vacuo and purified using HPLC (10-100% MeCN + 0.1% TFA, 0 –20 mins, 1.5 mL/min, $R_t$ = 10.6 min) to give the product as a pink/purple solid, TFA salt (3.9 mg, 29%). $^1$H NMR (700 MHz, CD3CN) δ 8.53 (d, J = 6.1 Hz, 1H), 8.31 (t, J = 7.7 Hz, 1H), 7.85 (d, J = 6.6 Hz, 1H), 7.79 – 7.75 (m, 1H), 7.70 (pd, J = 7.6, 1.5 Hz, 2H), 7.60 – 7.57 (m, 1H), 7.40 – 7.37 (m, 1H), 7.13 (d, J = 9.2 Hz, 2H), 6.81 (s, 1H), 6.53 (dd, J = 9.2, 2.1 Hz, 2H), 6.40 (d, J = 2.1 Hz, 2H), 4.26 (t, J = 7.6 Hz, 8H), 3.54 (d, J = 6.0 Hz, 2H), 3.34 – 3.29 (m, 3H), 3.25 (d, J = 11.8 Hz, 4H), 3.21 (t, J = 6.6 Hz, 4H), 2.49 (p, J = 7.6 Hz, 4H), 2.44 (t, J = 6.4 Hz, 2H), 2.27 (t, J = 6.3 Hz, 2H). MS (ESI): Calcd for $C_{42}H_{46}N_6O_4^{2+}$ $[M]^{2+}$: 349.18, found: 349.2.

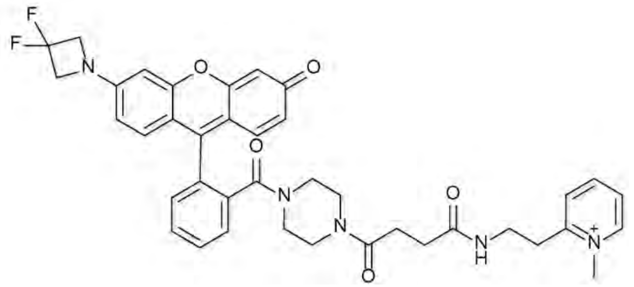

***5a, 503MP***: JF503 (13.6 mg, 0.033 mmol, 1.0 equiv) was added to a flame-dried reaction flask in 1.5 mL dry DMF. DIPEA (36 uL, 0.2 mmol, 6.3 equiv) was added. The reaction mixture was cooled to 0°C and HATU (30 mg, 0.079 mmol, 2.4 equiv) was added as a solid. The reaction mixture was stirred for 15 minutes at 0°C. Deprotected piperazine methylpyridinium linker, **3** (16.3 mg, 0.053 mmol, 1.6 equiv) was dissolved in 0.4 mL dry DMF and added to the reaction mixture and stirred overnight at RT. The reaction was concentrated in vacuo and purified using HPLC (10-100% MeCN + 0.1% TFA, 0–20 mins, 1.5 mL/min, $R_t$ = 13.2 min) to give the product as an orange solid, TFA salt (4.8 mg, 18.7%). $^1$H NMR (700 MHz, MeOD) δ 8.84 (d, J = 5.5 Hz, 1H), 8.42 (t, J = 7.7 Hz, 1H), 7.98 (d, J = 7.8 Hz, 1H), 7.88 (s, 1H), 7.84 – 7.78 (m, 2H), 7.77 – 7.72 (m, 1H), 7.56 (dd, J = 6.2, 2.3 Hz, 1H), 7.46 (d, J = 9.3 Hz, 2H), 7.15 – 7.11 (m, 1H), 7.06 (d, J = 8.8 Hz, 1H), 6.93 (d, J = 9.2 Hz, 1H), 6.90 (d, J = 2.1 Hz, 1H), 4.81 (t, J = 11.4 Hz, 4H), 2.56 (s, 2H), 2.40 (s, 2H). HRMS (ESI): Calcd for $C_{39}H_{38}F_2N_5O_5^+$ [M]$^+$: 694.28, found: 694.2829.

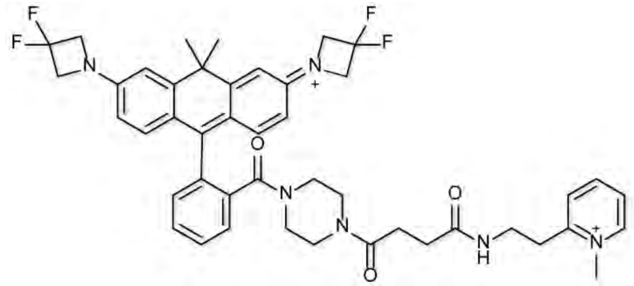

***6a, 585MP***: To JF585 (0.0137 g, 0.027 mmol, 1 equiv) dissolved in DCM (900 μL), a catalytic amount of DMF (6 μL) was added. Then oxalyl chloride (0.017 g, 0.13 mmol, 5 equiv, 0.012 mL) was added dropwise and the reaction was stirred at RT for 1 hr. Afterwards, TEA (0.016 g, 0.16 mmol, 6 equiv, 0.022 mL) and the deprotected piperazine methylpyridinium linker, **3** (0.0225 g, 0.054 mmol, 2 equiv) dissolved in DMF (150 μL) was added. The reaction was stirred at RT for 3 hrs. The reaction mixture was concentrated in vacuo and was purified using HPLC (10-100% MeCN + 0.1% TFA 0–20 min, 1.5 mL/min, $R_t$= 15.4 min) to give the product as a blue solid, TFA salt (0.0056 g, 26%). $^1$H NMR (700 MHz, MeOD) δ 8.86 – 8.83 (d, J = 5.9 Hz, 1H), 8.45 – 8.41 (t, J = 7.8 Hz, 1H), 8.00 – 7.96 (d, J = 7.6 Hz, 1H), 7.91 – 7.87 (s, 1H), 7.76 – 7.73 (dt, J = 7.6, 3.8 Hz, 2H), 7.66 – 7.63 (m, 1H), 7.48 – 7.44 (dd, J = 5.3, 3.2 Hz, 1H), 7.29 – 7.24 (d, J = 9.1 Hz, 2H), 7.05 – 7.02 (d, J = 2.1 Hz, 2H), 6.62 – 6.58 (dd, J = 9.1, 1.9 Hz, 2H), 4.82 – 4.73 (t, J = 11.2 Hz, 8H), 4.40 – 4.36 (s, 3H), 3.66 – 3.60 (s, 2H), 3.42 – 3.36 (d, J = 18.2 Hz, 4H), 3.35 – 3.32 (d, J = 6.6 Hz, 3H), 2.57 – 2.53 (t, J = 6.2 Hz, 2H), 2.42 – 2.38 (t, J = 6.3 Hz, 2H), 1.88 – 1.84 (s, 3H), 1.69 – 1.66 (s, 3H). HRMS (ESI) Calcd for $C_{45}H_{48}F_4N_6O_3^{2+}$ [M]$^{2+}$: 398.18, found: 398.1855.

4a, 549MP

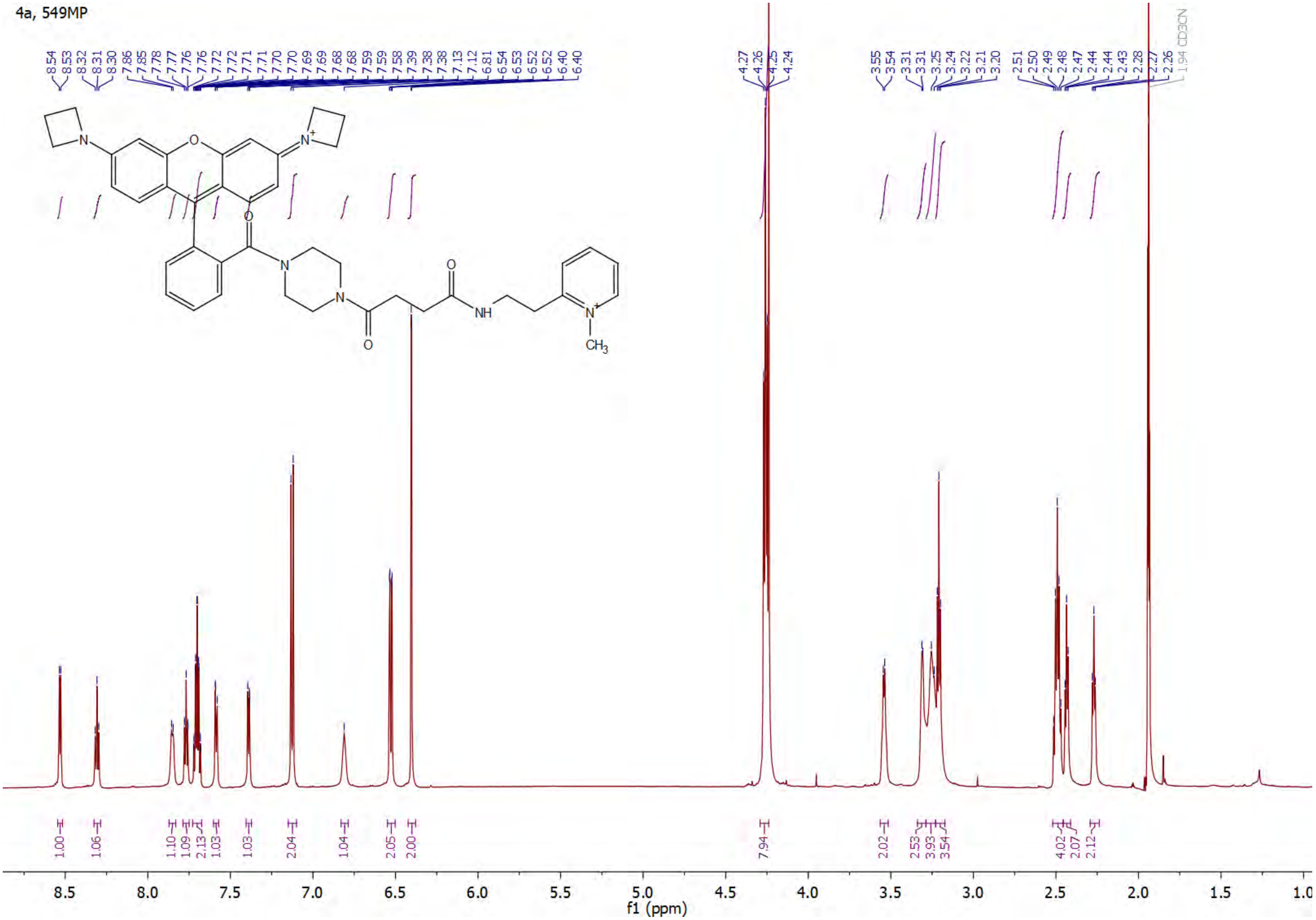

4a, 549MP

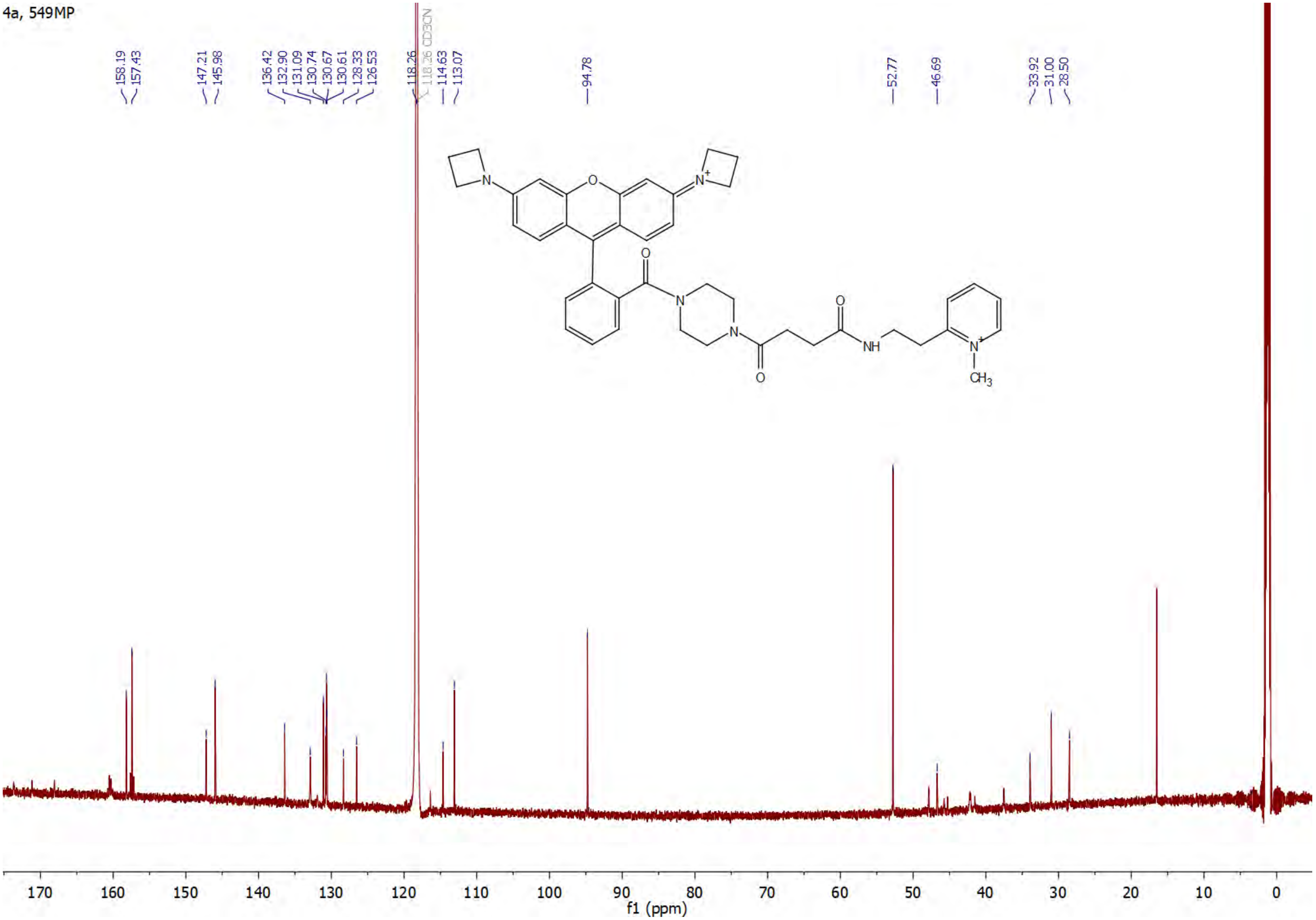
158.19
157.43
147.21
145.98
136.42
132.90
131.09
130.74
130.67
130.61
128.33
126.53
118.26
118.26 CD3CN
114.63
113.07
94.78
52.77
46.69
33.92
31.00
28.50
NH
CH3
170
160
150
140
130
120
110
100
90
80
70
60
50
40
30
20
10
0
f1 (ppm)

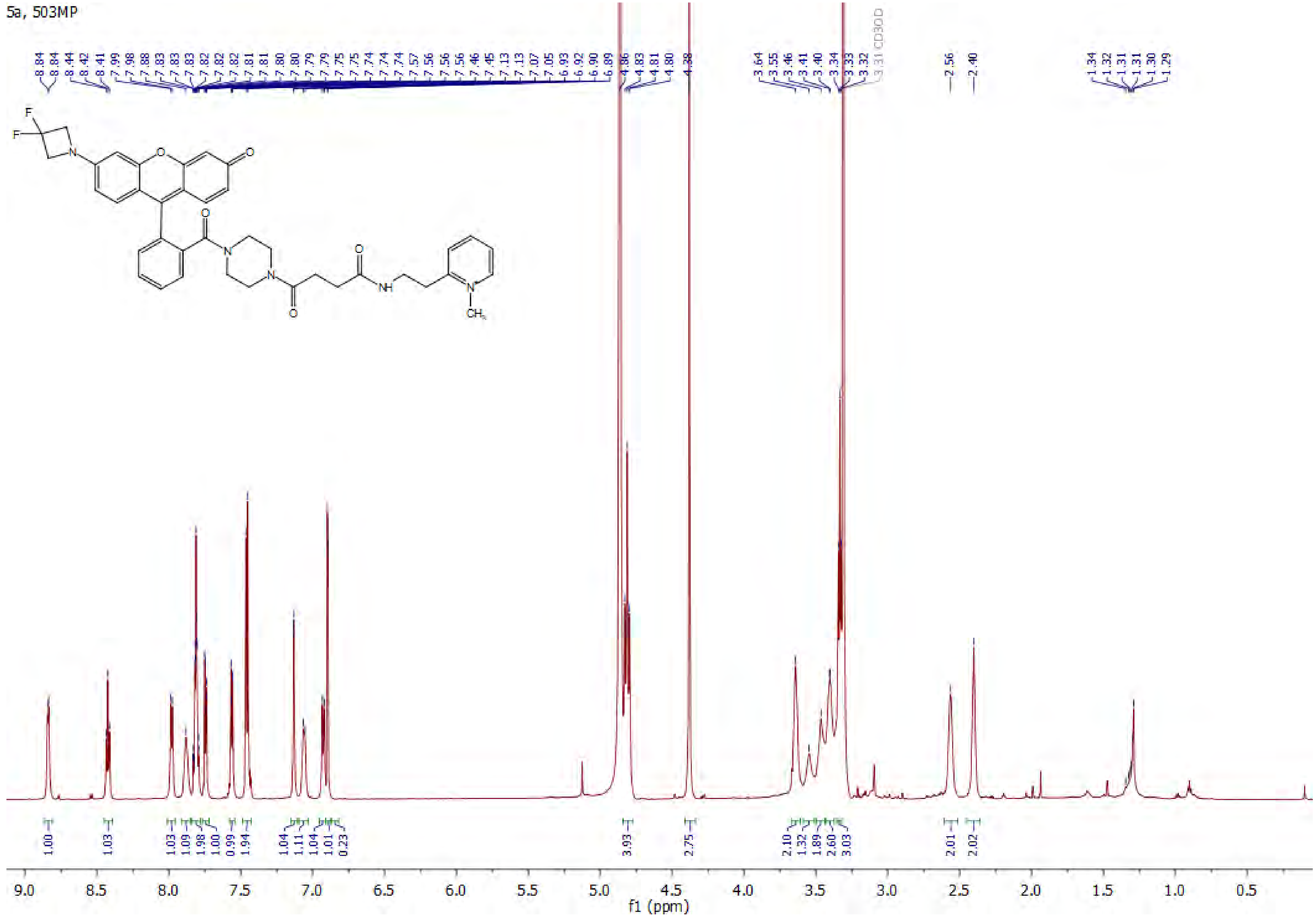
5a, 503MP
CD3OD
f1 (ppm)

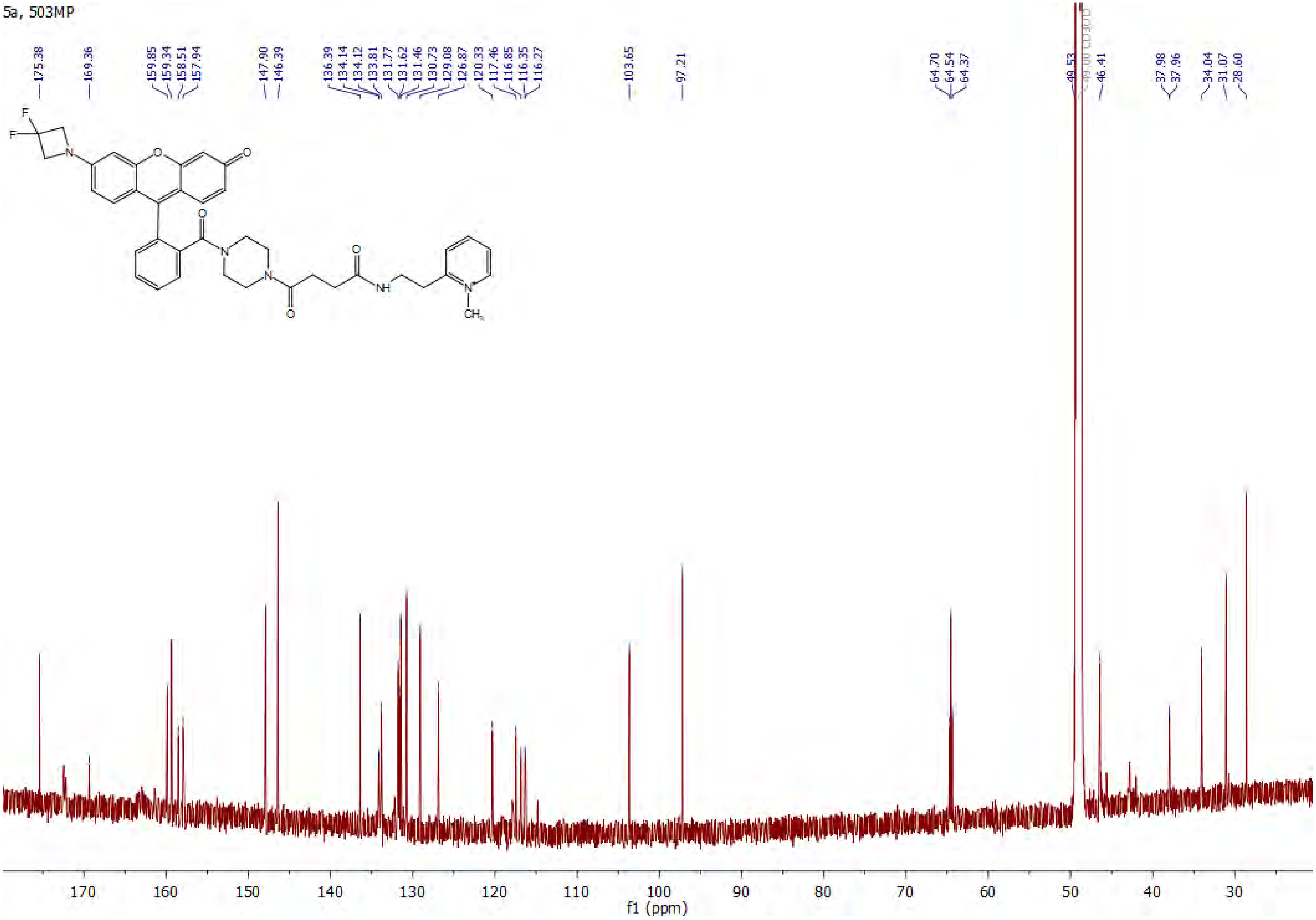
5a, 503MP
175.38
169.36
159.85
159.34
158.51
157.94
147.90
146.39
136.39
134.14
134.12
133.81
131.77
131.62
131.46
130.73
129.08
126.87
120.33
117.46
116.85
116.35
116.27
103.65
97.21
64.70
64.54
64.37
49.53
46.41
37.98
37.96
34.04
31.07
28.60
F
F
N
O
O
O
N
N
O
O
NH
N+
CH3
170 160 150 140 130 120 110 100 90 80 70 60 50 40 30
f1 (ppm)

6a, 585MP

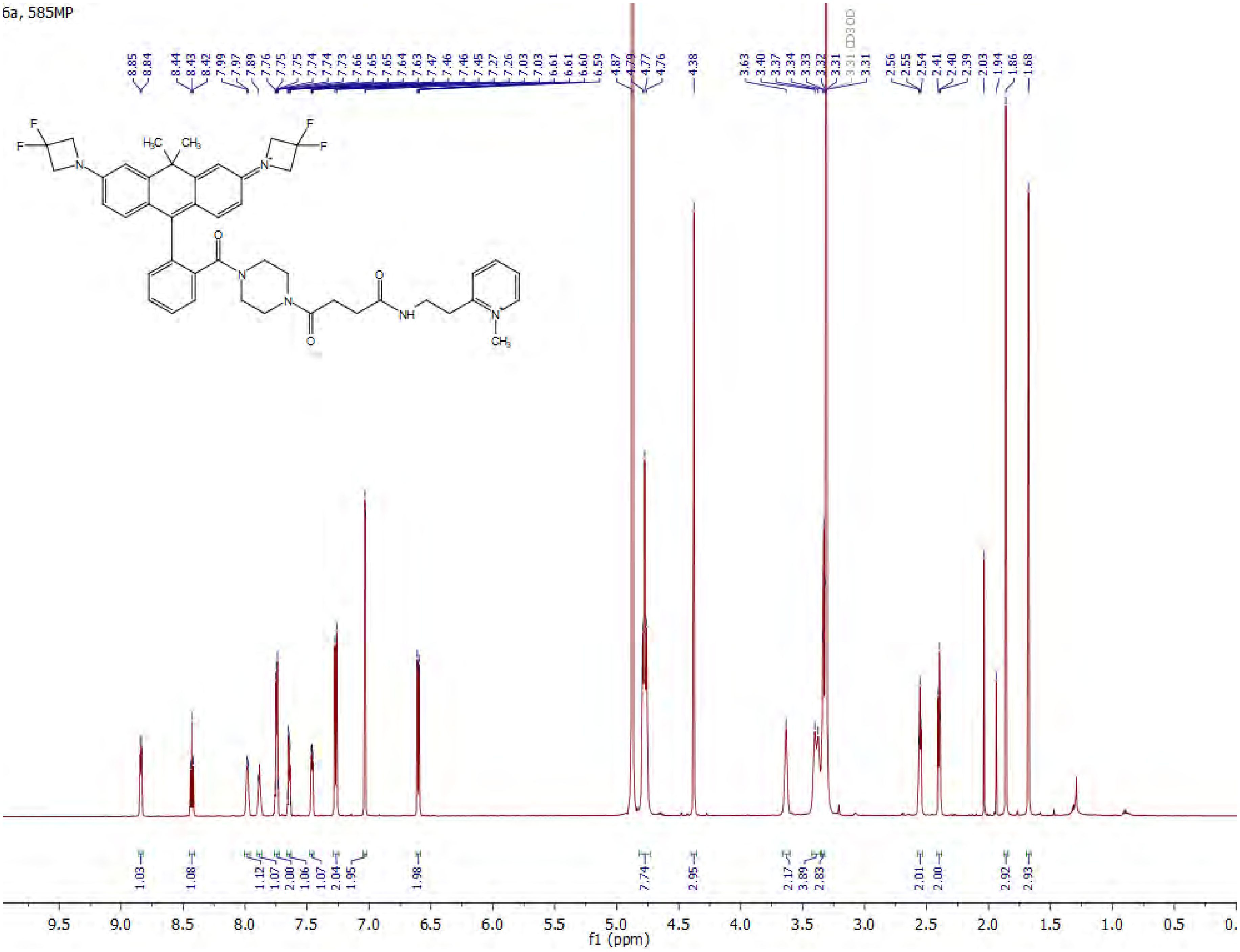

6a, 585MP

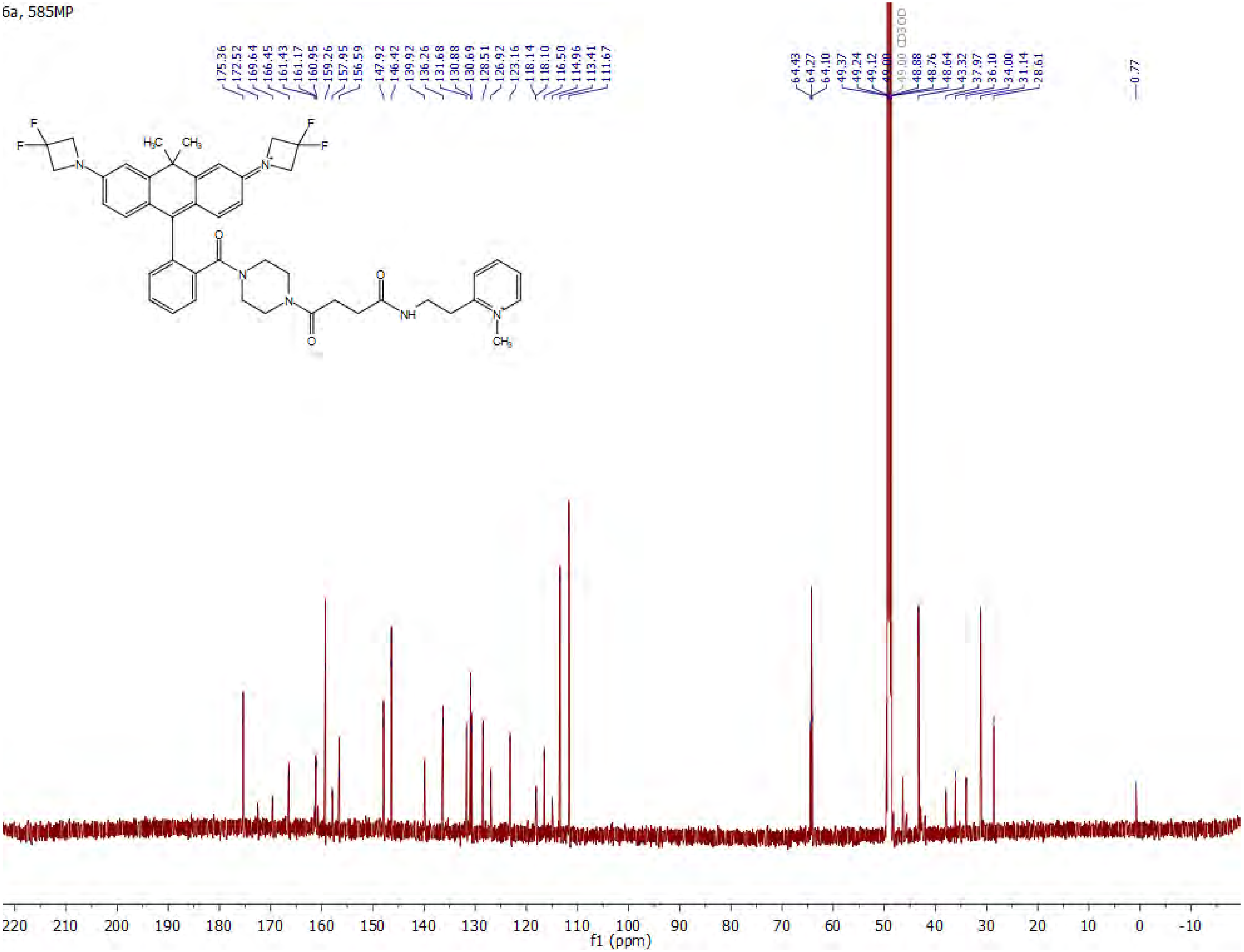

1. Clark Still, W., Kahn, M. & Mitra, A. Rapid Chromatographic Technique for Preparative Separations with Moderate Resolution. *Journal of Organic Chemistry* **43**, 2923–2925 (1978).
2. Nissen, J. C. & Tsirka, S. E. Tuftsin-driven experimental autoimmune encephalomyelitis recovery requires neuropilin-1. *Glia* **64**, 923–936 (2016).
3. Dunn, K. W., Kamocka, M. M. & Mcdonald, J. H. A practical guide to evaluating colocalization in biological microscopy. *Am J Physiol Cell Physiol* **300**, 723–742 (2011).
4. Grimm, J. B. *et al.* A general method to improve fluorophores for live-cell and single-molecule microscopy. *Nat Methods* **12**, 244 (2015).
5. Grimm, J. B. *et al.* A general method to fine-tune fluorophores for live-cell and in vivo imaging. *Nat Methods* **14**, 987 (2017).
6. Grimm, J. B. *et al.* Carbofluoresceins and carborhodamines as scaffolds for high-contrast fluorogenic probes. *ACS Chem Biol* **8**, 1303–1310 (2013).
7. Preston, A. N. *et al.* Visualizing the Brain's Astrocytes with Diverse Chemical Scaffolds. *ACS Chem Biol* **13**, 1493–1498 (2018).
8. Pierangelo Gobbo, Praveen Gunawardene, Wilson Luo & Mark S. Workentin. Synthesis of a Toolbox of Clickable Rhodamine B Derivatives. Synlett, 26(09), 1169–1174 | 10.1055/s-0034-1380191. *Synlett* **26**, 1169–1174 (2015).
